# A Mechanically Weak Extracellular Membrane-Adjacent Domain Induces Dimerization of Protocadherin-15

**DOI:** 10.1101/460097

**Authors:** P. De-la-Torre, D. Choudhary, R. Araya-Secchi, Y. Narui, M. Sotomayor

## Abstract

The cadherin superfamily of proteins is defined by the presence of extracellular cadherin (EC) repeats that engage in protein-protein interactions to mediate cell-cell adhesion, cell signaling, and mechanotransduction. The extracellular domains of non-classical cadherins often have a large number of EC repeats along with other subdomains of various folds. Protocadherin-15 (PCDH15), a protein component of the inner-ear tip link filament essential for mechanotransduction, has eleven EC repeats and a membrane adjacent domain (MAD12) of atypical fold. Here we report the crystal structure of a pig PCDH15 fragment including EC10, EC11, and MAD12 in a parallel dimeric arrangement. MAD12 has a unique molecular architecture and folds as a ferredoxin-like domain similar to that found in the nucleoporin protein Nup54. Analytical ultracentrifugation experiments along with size exclusion chromatography coupled to multi-angle laser light scattering and small-angle X-ray scattering corroborate the crystallographic dimer and show that MAD12 induces parallel dimerization of PCDH15 near its membrane insertion point. In addition, steered molecular dynamics simulations suggest that MAD12 is mechanically weak and may unfold before tip-link rupture. Sequence analyses and structural modeling predict the existence of similar domains in cadherin-23, protocadherin-24, and the “giant” FAT and CELSR cadherins, indicating that some of them may also exhibit MAD-induced parallel dimerization.

## INTRODUCTION

The cadherin superfamily of proteins can be broadly divided into three main subfamilies that include the classical cadherins (type I, II, and desmosomal), the clustered protocadherins (α, β, and γ), and the non-classical non-clustered cadherins (δ1 and δ2 protocadherins along with various other atypical cadherins) (1–4). All of them feature two or more tandem extracellular cadherin (EC) “repeats” of similar fold (Greek key motifs with seven β strands) but distinct residue sequence (5–8). These EC repeats are often involved in protein-protein interactions essential for the diverse functions played by cadherins in cell-cell adhesion (2,9,10), signaling (11–15), and mechanotransduction (16–18).

The vertebrate classical cadherins were discovered as glycoproteins involved in calcium (Ca^2+^)-dependent cellular adhesion (19,20), and the mechanism by which they facilitate adhesion is well understood (2). Their processed extracellular domains (21), with five EC repeats, protrude from the surface of adjacent cells to engage in mostly homophilic (same type) binding across the cells (*trans*) (2). The individual *trans* bonds linking adjacent cells are weak, with the strength of cell-cell adhesion stemming from multiple *trans* bonds supported by *cis* interactions between extracellular domains protruding from the same cell (22,23). Biophysical and structural studies of classical cadherins suggest that their *trans* adhesive interactions are mediated by EC1 to EC1 contacts, while *cis* interactions of classical type I cadherins involve EC1 of one molecule interacting with EC2 of the neighboring molecule (23–31). In contrast, clustered protocadherins, with six EC repeats, use an antiparallel binding interface involving repeats EC1 to EC4 for adhesion (32–36), with promiscuous *cis* interactions mediated by the EC repeats closest to the membrane (EC5 and EC6) (37). The m2 protocadherins seem to use a similar mechanism for *trans* interactions with EC1-4 antiparallel interfaces (38), yet it is unclear if the u protocadherins engage in clustered-protocadherin-like *cis* interactions.

Less is known about binding mechanisms and oligomerization states of non-classical, non-clustered atypical cadherins, especially those that have enormous extracellular domains, such as protocadherin-15 (PCDH15) and cadherin-23 (CDH23), protocadherin-24 (PCDH24), the FAT cadherins, and CELSRs. The PCDH15 and CDH23 proteins, involved in hereditary deafness and blindness (18,39–41), use their EC1-2 tips to engage in an antiparallel, heterophilic *trans*“handshake” interaction essential for inner-ear function (42–44). Electron microscopy of negatively stained extracellular domains also suggests that their full-length extracellular domains form parallel *cis* homodimers (42). Although most PCDH15 and CDH23 fragments with up to four EC repeats have failed to conclusively show bona-fide homophilic *cis* interactions (45–47), two recent structures show that PCDH15 can dimerize at EC2-3 (48) and at its C-terminal domain near the membrane (49). Less is known about the specific contacts leading to parallel dimerization of CDH23. Similarly, PCDH24 forms *trans* heterophilic bonds with the mucin-like cadherin CDHR5, and also forms *trans* homophilic bonds with itself (50). The PCDH24-CDHR5 interactions are essential for inter-microvillar link formation and important for brush-border function (51), but little is known about the EC repeats involved or about PCDH24’s oligomerization states. Further, the giant cadherins FAT4 and DCHS1 have been shown to self bend and form heterophilic complexes involved in cell signaling, with repeats EC1 to EC4 being sufficient for the interaction (52), yet details are still missing. The extracellular domains of FAT1-3 (13) and CELSR1-3 (53) have been the least studied biochemically, and their binding modes and oligomerization states remain to be fully elucidated.

Interestingly, long non-classical, non-clustered atypical cadherins feature extracellular domains with modules that do not fold as canonical cadherin repeats. PCDH15 and CDH23 have membrane adjacent domains (MADs) that are ~100-residues long and that have no predicted fold (Fig. 1 *A*). For PCDH15, this domain has also been labeled as “extracellular linker” (EL) (49) or “protocadherin-15 interacting-channel associated domain” (PICA) (48). PCDH24 has a similar putative MAD of unknown fold, while FAT and CELSR cadherins all have “unknown” domains between their cadherin repeats and additional extracellular domains with predicted EGF, EGF-like or LamG folds (details below). Here we present an X-ray crystal structure of the pig (*Sus scrofa*[*ss*]) PCDH15 including repeats EC10, EC11, and its MAD12 fragment (Fig. 1 *B* and Fig. S1). This structure reveals a ferredoxin-like fold (49) for MAD12 that induces parallel *cis* homodimerization, that is predicted to be mechanically weak using steered molecular dynamics (SMD) simulations, and that might be present in CDH23, PCDH24 and the FAT and CELSR cadherins, as suggested by sequence analyses and structural modeling.

**FIGURE 1.**
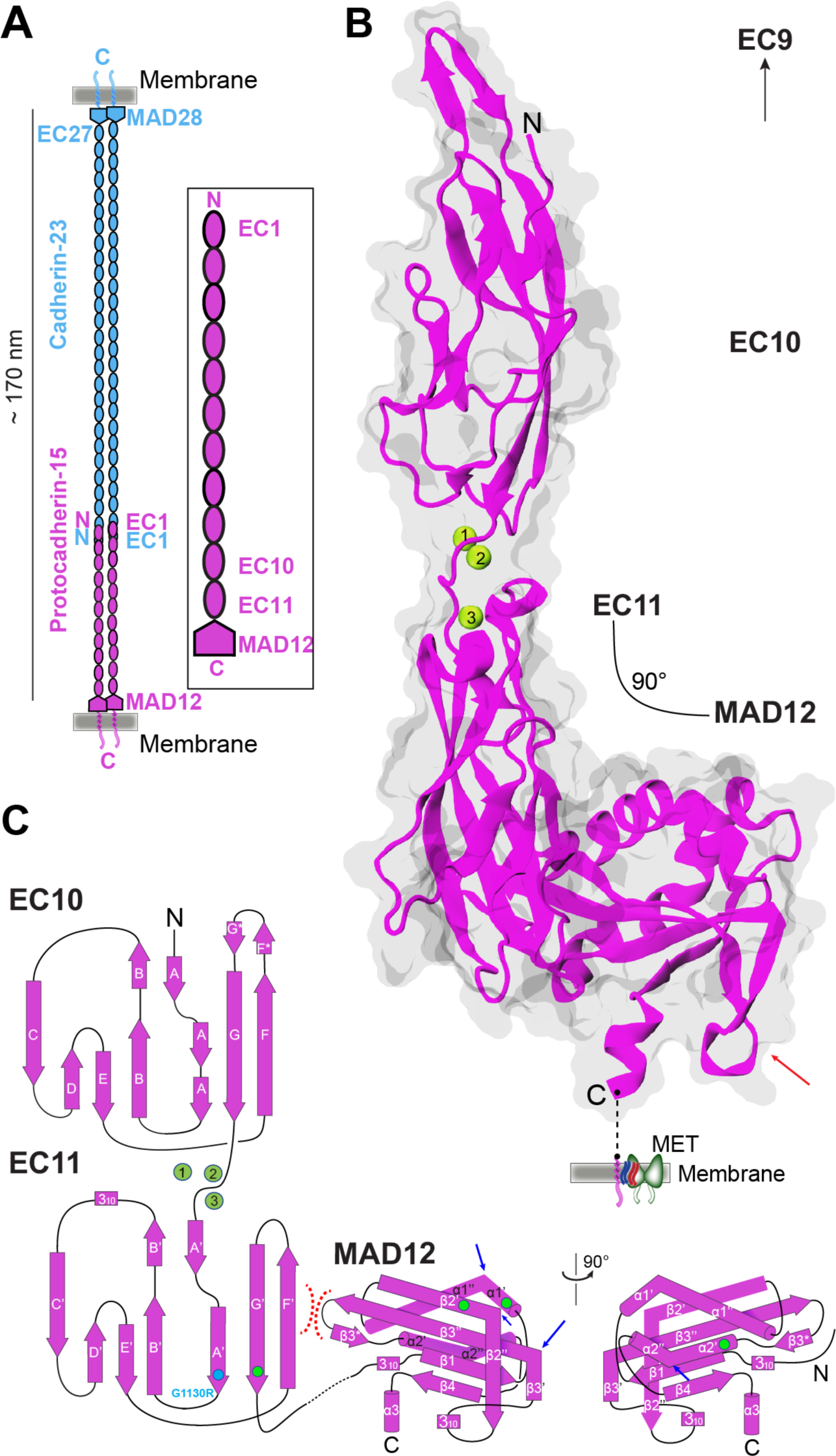
PCDH15 architecture and structure of the EC10-MAD12 fragment. (**A**) Schematic of the cadherin tip link and the PCDH15 extracellular domain. Inset shows the location of the fragment studied here. (**B**) Ribbon diagram of *ss* PCDH15 EC10-MAD12 (magenta) with transparent molecular surface. Ca^2+^ ions are 5 shown as green spheres. Red arrow indicates β”-3_10_-β3’ hook. Dashed line indicates unresolved region. Membrane and mechanotransduction channel (MET) are indicated (not to scale). (**C**) Topology diagram of PCDH15 EC10-MAD12. A typical cadherin fold is observed for EC10 and EC11. A ferredoxin-like fold is observed for MAD12. Intramolecular EC11-MAD12 contact is highlighted by red dashed lines. Blue arrows indicate kinks. Cyan and green circles highlight sites involved in hereditary deafness (causal and correlated 10 mutations, respectively).

## MATERIALS AND METHODS

### Cloning, Expression, and Purification of Bacterially Expressed PCDH15 EC10-MAD12

Pig and mouse PCDH15 including EC10, EC11, and MAD12 (*ss* and *mm* PCDH15 EC10-MAD12) (Tables S1 and S2) were subcloned into NdeI and XhoI sites of the pET21a vector. These constructs were used for expression in Rosetta (DE3) competent cells (Novagen) cultured in TB and induced at OD _600_ = 0.6 with 1 mM of IPTG at 30 °C for ~16 hr. Cells were lysed by sonication in denaturing buffer (20 mM TrisHCl [pH 7.5], 6 M guanidine hydrochloride, 10 mM CaCl_2_, and 20 mM imidazole). The cleared lysates were incubated with Ni-Sepharose (GE Healthcare) for 1 h and eluted with denaturing buffer supplemented with 500 mM imidazole. Both pig and mouse PCDH15 EC10-MAD12 proteins were refolded at 4 °C by fast or drop-wise dilution of 20 ml of eluted denatured protein (1-2 mg/ml) into 480 ml of refolding buffer (20 mM TrisHCl [pH 8.0], 150 mM KCl, 5 mM CaCl_2_, 400 mM L-arginine, 2 mM DTT or 1 mM TCEP, and 10% glycerol). For fast dilution refolding, the denatured protein was dispensed into the refolding buffer over a 1-min period with vigorous stirring. For drop-wise refolding, the denatured protein was dispensed over a 20-min period with gentle stirring. In both cases, the protein solution was kept under constant and gentle stirring after mixing. All protein solutions were concentrated to 10 ml using Amicon Ultra-15 filters at 3,000 rpm with mixing every 20 min. Refolded protein was further purified on a Superdex-200 column (GE Healthcare) in holding buffer containing 20 mM TrisHCl [pH 8.0], 150 mM KCl, 5 mM CaCl_2,_ and 1 mM TCEP. Purity of the recombinant protein was analyzed by SDS-PAGE, after which the protein-containing fractions were pooled and used for further experiments. Pure proteins were concentrated by centrifuge ultrafiltration (Vivaspin 6 10 kDa) to 3-5 mg/ml for crystallization trials and biochemical assays. Protein concentrations were determined using the NanoDrop (Thermo Scientific).

### Cloning, Expression, and Purification of Mammalian-Expressed *mm\** PCDH15 EC10-MAD12

Suspension Expi293F cells (Thermo Fisher) were cultured in Expi293 medium at 37 °C, 8% CO_2_, and incubated on an orbital shaker at 125 rpm. The coding sequence for *mm* PCDH15 EC10-MAD12 was inserted into a pHis-N1 expression vector (modified version of the pEGFP-N1 vector from Clontech where the EGFP has been substituted for a hexahistidine tag; James Jontes, The Ohio State University) using XhoI and KpnI restriction sites. The native signal sequence was included before EC10. For a typical 30 ml expression, cells were prepared at a density of 2.9 × 10^6^ cells/ml and exchanged into 25.5 ml of fresh media. The DNA complexes were prepared by diluting 30 µg of DNA up to 1.5 ml with Opti-MEM and diluting 81 µl of ExpiFectamine 293 up to 1.5 ml. After 5 minutes, the two solutions were combined, gently mixed, and incubated for an additional 20 min at room temperature. The complexes were then added to the cells and incubated for 20 hours. The next day, 150 µl of ExpiFectamine 293 Transfection Enhancer 1 and 1.5 ml of ExpiFectamine 293 Transfection Enhancer 2 were added. Cells were grown for 4 days, and the conditioned media (CM) was collected. The CM was dialyzed overnight against 20 mM TrisHCl [pH 7.5], 150 mM KCl, 50 mM NaCl, and 10 mM CaCl_2_ to remove EDTA. The CM was concentrated using Amicon Ultra-15 10 kD or 30 kD concentrators and incubated with Ni-Sepharose beads for 1 h. The beads were washed 3 times with 20 mM TrisHCl [pH 8.0], 300 mM NaCl, 10 mM CaCl_2_, and 20 mM imidazole, and the target protein was eluted with the same buffer containing 500 mM imidazole. The protein was further purified on a Superdex-200 column (GE Healthcare) in 20 mM TrisHCl [pH 7.5], 150 mM KCl, 50 mM NaCl, and 2 mM CaCl_2_. The protein was concentrated and the final concentration was determined by measuring absorbance using the NanoDrop (Thermo Scientific).

### Crystallization and Micro-seeding

Initial needle-like crystals of *ss* PCDH15 EC10-MAD12 were grown by vapor diffusion at 4 °C by mixing equal volumes (0.6 μl) of protein (purified without TCEP) and reservoir solutions (0.1 M HEPES [pH 7.5], 25% W/v PEG 2000 MME) and by equilibrating against 75 μl of reservoir solution in a sitting drop setup. A seed stock was prepared by mixing of crystal-containing solution (2 μl) with reservoir solution (50 μl) followed by vortexing with microseed beads (Molecular Dimensions MD2-14). The resulting solution was diluted to prepare a seed stock and used for random micro-seeding (54). Crystallization droplets were composed of equal volumes (1 μl) of protein solution (repurified without TCEP; 4.7 mg/ml) and reservoir solution along with 0.5 μl of the seed stock. The droplets were equilibrated against 75 μl of reservoir solution in a sitting-drop setup at 4 °C. Final hits were obtained in 0.1 M HEPES [pH 7.5], and 20% V/v jeffamine M-600.

### Data Collection and Structure Determination

Crystals were cryoprotected in reservoir solution plus 25% V/v PEG400. All crystals were cryo-cooled in liquid N_2_. X-ray diffraction data was collected as indicated in Table 1 and processed with HKL2000 (55). The final structure was determined by molecular replacement using PHASER (56) and Buccaneer (57). Initial search models were based on the structure of *hs* PCDH15 EC10 (PDB ID: 4XHZ) (45) and a separate homology model for the *mm* PCDH15 EC11 repeat obtained using SWISS-MODEL (58) and the PDB ID: 5DZX (32) as a template. MAD12 was built *de novo* and refined using COOT (59). REFMAC5 (60) was used for restrained TLS refinement. Data collection and refinement statistics are provided in Table 1. Coordinates for *ss* PCDH15 EC10-MAD12 have been deposited in the Protein Data Bank with entry code 6BXZ.

**TABLE 1.**
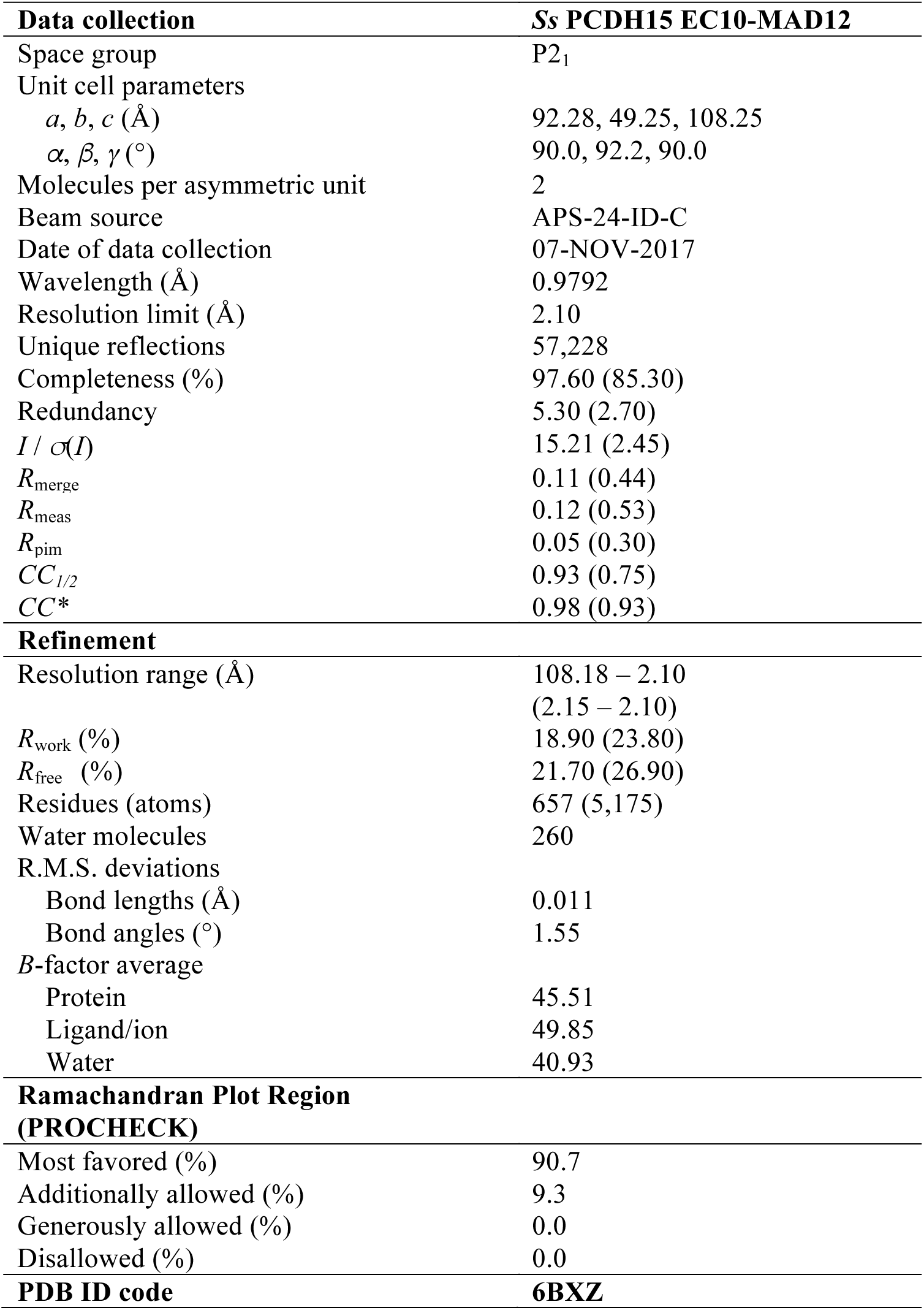
Statistics for *ss* PCDH15 EC10-MAD12 Structure.

### Analytical Ultracentrifugation

Sedimentation velocity AUC experiments were performed in a ProteomeLab XL-I analytical ultracentrifuge (Beckman Coulter) following standard procedures (61–63). Briefly, SEC purified protein samples were loaded into AUC cell assemblies with Epon centerpieces and 12 mm path length. To achieve chemical and thermal equilibrium, the An-50 TI rotor with loaded samples was allowed to equilibrate for ~2 h at 20 °C in the centrifuge. The rotor was spun at 50,000 rpm and data was collected using absorption optics. Data analysis was performed with the software SEDFIT (http://sedfitsedphat.nibib.nih.gov), using a continuous sedimentation coefficient distribution model *c*(*S*). Standard values for buffer viscosity (0.01002 poise), density (1 g/ml) and partial specific volume (0.73 ml/g) were used, and confidence level was set to 0.68 as routinely done. The obtained *c*(*S*) distribution was loaded in GUSSI (64). AUC experiments were done with at least two biological replicates, which were also accompanied by at least one duplicate to ensure data was obtained accurately.

### SEC-MALS

SEC-MALS data was collected using an ÄKTAmicro system connected in series with a Wyatt miniDAWN TREOS system. The protein samples were separated on a Superdex-S200 3.2/30 column (GE Healthcare), and both the absorbance at 280 nm and the light scattering were monitored. The scattering information was subsequently converted into molecular weight using a rod-like model. SEC-MALS experiments were done with at least two biological replicates and accompanied by at least one duplicate.

### SEC-SAXS

SAXS data were collected at SIBYLS beamline 12.3.1 in the Advanced Light Source (ALS) (Berkeley, CA) as described (65,66) (Table S3). Purified samples of *ss* PCDH15 EC10-MAD12 (6.3 mg/ml) and *mm* PCDH15 EC10-MAD12 (5.3 mg/ml) were analyzed. Data were collected using an Agilent 1260 series HPLC with a Shodex KW-803 analytical column at a flow rate of 0.5 ml/min using 20 mM TrisHCl [pH 8.0], 150 mM KCl, 5 mM CaCl2, and 1 mM TCEP as a mobile phase at 20 °C. Each 50 Cl sample was run through SEC and 3 s X-ray exposures were collected continuously during a ~40 min elution. Samples were examined with incident light at λ = 1.03 Å and at a sample to detector distance of 1.5 m resulting in scattering vectors, *q*, ranging from 0.01 Å^−1^ to 0.6 Å^−1^, where the scattering vector is defined as *q*= 4π sinθ / λ and 2θ is the measured scattering angle. SAXS frames recorded prior to the protein elution peak were used to subtract all other frames. Buffer subtraction and data reduction was performed at the beamline with SCÅTTER (67). Buffer matched controls were used for buffer subtraction.

Further data analysis of the merged SAXS data was carried out with PRIMUS (68) and the ATSAS program suite (69). Estimates of the radius of gyration (*R*_g_) from the Guinier region were measured with PRIMUS. Maximum dimension (*D*_max_) of particles was estimated from an indirect Fourier transform of the SAXS profiles using GNOM (70). Values of *D*_max_ between 120 and 140 Å provided the best solutions for both samples. The oligomeric state of the samples was assessed by estimating their molecular weight using the method implemented in the SAXSMoW2 server (71). Three-dimensional spatial representations of *ss* and *mm* PCDH15 EC10-MAD12, classified as flat using DATCLASS and with an AMBIMETER shape ambiguity score of 2.3 for both samples (possibly ambiguous) (72), were obtained by *ab initio* modeling with DAMMIF (73) considering *q* < 0.2, P1 symmetry, and prolate anisometry. For each sample, thirty models were generated and averaged with DAMAVER (74) with a mean normalized spatial discrepancy (NSD) of 1.07 ± 0.16 (one model rejected) and 0.98 ± 0.13 (two models rejected) for *ss* and *mm* PCDH15 EC10-MAD12, respectively. A final bead model refined against the SAXS data was produced with DAMMIN/DAMMSTART (75) with an estimated resolution of 44 ± 3 Å and 37 ± 3 Å (76). Model scattering intensities were computed from *ss* PCDH15 EC10-MAD12 (6BXZ) and fitted to the experimental SAXS data using FoXS (77). Search of conformational changes of the dimeric structure in solution was performed by normal modes analysis with SREFLEX (78).

### Molecular Dynamics Simulations

The structure of *ss* PCDH15 EC10-MAD12 (PDB ID 6BXZ) was used to build monomeric and dimeric systems. Missing residues in one of the dimer subunits were included in the structure using coordinates obtained from the other superposed subunit. Final models included residues p.G1009 to p.I1342. The psfgen, solvate, and autoionize VMD plugins were used to build all systems (Table S4) (79). Hydrogen atoms were automatically added to protein and crystallographic water molecules. Residues D, E, K and R were charged. Histidine residues were neutral, and their protonation states were chosen to favor the formation of evident hydrogen bonds. Additional water molecules and randomly placed ions were used to solvate the systems (150 mM KCl) in boxes large enough to accommodate stretched states and to prevent interactions between periodic images.

MD simulations were performed using NAMD 2.12 (80), the CHARMM36 force field for proteins with the CMAP correction and the TIP3P model for water (81). A cutoff of 12 Å (with a switching function starting at 10 Å) was used for van der Waals interactions along with periodic boundary conditions. The Particle Mesh Ewald method was used to compute long-range electrostatic forces (grid point density of >1 Å^−3^). A uniform 2 fs integration time step was used together with SHAKE. Langevin dynamics was utilized to enforce constant temperature (*T* = 300 K; γ = 0.1 ps^−1^). Constant pressure simulations (*NpT*) at 1 atm were done using the hybrid Nosé-Hoover Langevin piston method with a 200 fs decay period and a 100 fs damping time constant.

Constant-velocity stretching simulations used the SMD method and the NAMD Tcl forces interface. Constant-velocity SMD simulations (82–85) were performed by attaching Cα atoms of N- and C-terminal residues to independent virtual springs. For some simulations (Table S4), these springs were connected to virtual slabs connected to a third spring. All springs had stiffness *k*_*s*_ = 1 kcal mol^−1^ Å^−2^. The free ends of the stretching springs were moved away from the protein in opposite directions at a constant velocity. Applied forces were computed using the extension of the virtual springs. Maximum force peaks and their averages were computed from 50 ps running averages used to eliminate local fluctuations.

### Sequence Analyses, Structural Modeling, and Analysis Tools

To compare protein sequences among various species, 18 sequences from 17 species were obtained for their longest PCDH15 isoforms from the NCBI protein database (Table S1) and were split into the corresponding EC10, EC11, and MAD12 fragments before alignment. These fragments were aligned as in (47). The ClustalW algorithm (86) on Geneious (87) was used to obtain percent sequence identity for each fragment. Alignment files from Geneious were colored according to % of sequence identity in JalView with 45% conservation threshold (88). Similarly, sequences among MADs from different cadherin subfamilies (Table S2) were split for sequence alignments (colored according to % sequence identity in JalView with 1% conservation threshold), which were used as input into the Sequence Identity and Similarity (SIAS) server (89) to obtain sequence identity matrices. Residue numbering throughout the text and in the structures corresponds to the processed protein, which does not include the signal peptide. Numbering is based on entries *hs* PCDH15 isoform 1 precursor, NCBI Reference Sequence: NP_001136235.1 (signal peptide with 26 residues). Details of interatomic interactions were analyzed using Maestro (90) and VMD (79). Molecular stereo images were prepared with PyMOL (91). Plots and curve fits were prepared with XMGrace or QtiPlot. Protein-protein contacts were analyzed with the Protein, Interfaces, Surfaces, and Assemblies (PISA) server (92). Coordinates for structural models of *hs* CDH23 MAD28, *hs* PCDH24 MAD10, *hs* FAT1 MAD35, *hs* FAT2 MAD34, *hs* CELSR1 MAD10, *hs* CELSR2 MAD10, *dm* FAT MAD35, as well as dimeric *hs* CDH23 EC26-MAD28 and *hs* PCDH24 EC8-MAD10 are available upon request.

### Protein-Protein Docking and MM/GBSA Calculations

Homology models of *hs* CDH23 EC26-27 and *hs* PCDH24 EC8-9 were obtained using SWISS-MODEL (58) and published structures (PDB IDs 5TFK and 5SZR) (33,47) as templates. RaptorX MAD models were connected to models of the preceding EC repeats to build L-shaped structures using COOT (59) and *ss* PCDH15 EC10-MAD12 as a guide. Models were minimized to reduce steric clashes with Macromodel and the OPLS3e force field using standard parameters (implicit solvent, charges from forcefield, PRCG method with 2500 iterations in gradient conversion mode, convergence threshold of 0.05) without constraints (93). Resulting monomers were built as dimers based on *ss* PCDH15 EC10-MAD12 and further minimized (94). Monomeric protomers were then used for protein-protein docking with BioLuminate using the “Dimer mode” with 70,000 rotations for the ligand. Final docking poses were selected after automatized clustering and structural superposition to *ss* PCDH15 EC10-MAD12. Final models were further minimized and used as inputs for MM/GBSA calculations using Prime (93), the VSGB solvation model and the OPLS3e force field with standard parameters, the “side chain only” minimization mode, and no constraints.

## RESULTS AND DISCUSSION

Multiple sequence alignments of the PCDH15 EC10-MAD12 from 17 different species reveal that repeat EC10 is less conserved compared to repeat EC11, which is moderately conserved, while MAD12 is most conserved, with sequence identities of 37.6%, 47.7%, and 58.3%, respectively (Fig. S2). Secondary structure predictions and domain identification algorithms are consistent in identifying PCDH15 EC10 and EC11 as EC repeats, while failing to identify MAD12 as a domain of known fold. As expected, Ca^2+^-binding motifs for the linker region EC10-11, typically characterized by the sequence NTerm-XEX-DXD-D(R/Y)E-XDX-DXNDN-CTerm, are highly conserved, with a minor variation at the linker (DENXH). In contrast, Ca^2+^-binding motifs are absent in the EC11-MAD12 linker, despite the high sequence conservation of the last domain across mammals, birds, reptiles, and fish (Fig. S2). To determine the architecture and fold of this fragment and to biochemically characterize it, we purified *ss* and *Mus musculus*(*mm)* PCDH15 EC10-MAD12 from bacterial inclusion bodies, as well as *mm* PCDH15 EC10-MAD12 produced in mammalian cells (*mm*\*). These protein fragments included residues p.H1007 to p.E1353 (residue numbering corresponds to processed protein without its signal sequence). All protein fragments were well behaved in size exclusion chromatography (SEC) experiments (Fig. S3) and were used for crystallization trials, structure determination, and biochemical and biophysical analyses, as described below.

### Crystal Structure of Pig PCDH15 EC10-MAD12 Reveals a Ferredoxin-like Fold for MAD12

Diffracting crystals were obtained for the *ss* PCDH15 EC10-MAD12 protein fragment, for which an X-ray crystal structure was solved and refined to 2.1 Å resolution (Table 1). Two protomers were observed in the asymmetric unit, comprising residues p.G1009 to p.I1342 and p.I1011 to p.T1340, with missing residues not observed in the electron density map. These protomers adopted an L-shaped architecture (Fig. 1 *B*) and interacted with each other in a parallel *cis* arrangement, as also recently reported in (49) and as discussed below. The long end of the L-shaped, bent structure has two EC repeats (EC10 and EC11) with typical Greek key folds (seven β strands labeled A to G for EC10; A’ to G’ for EC11; Fig. 1 *C*) and a canonical linker region with three bound Ca^2+^ ions at sites 1, 2, and 3 (Figs. 1 *B* and 2 *A*; Fig. S1 *A*). The *ss* PCDH15 EC10 repeat shares structural features with its orthologs from human and mouse (45), with some differences in its FG connector, which folds into a stableff helix in the presence of the preceding EC9 repeat to be part of a bent, Ca^2+^-free EC9-10 linker region. Instead, in one chain of our structure, this loop is extended forming a β hairpin (F*G*) involved in crystal contacts (Fig. 1, *B* and *C*), while electron density is absent for the other chain. The *ss* PCDH15 EC11 repeat has a canonical fold with an atypical and highly conserved p.N1223-XL-p.D1226 linker that forms a 3_10_ helix leading to MAD12 (Fig. S2), a domain that is positioned next to the EC11 face formed by β-strands A’, F’, and G’. MAD12 tucks against EC11 to form the short arm of the L-shaped protomer.

**FIGURE 2.**
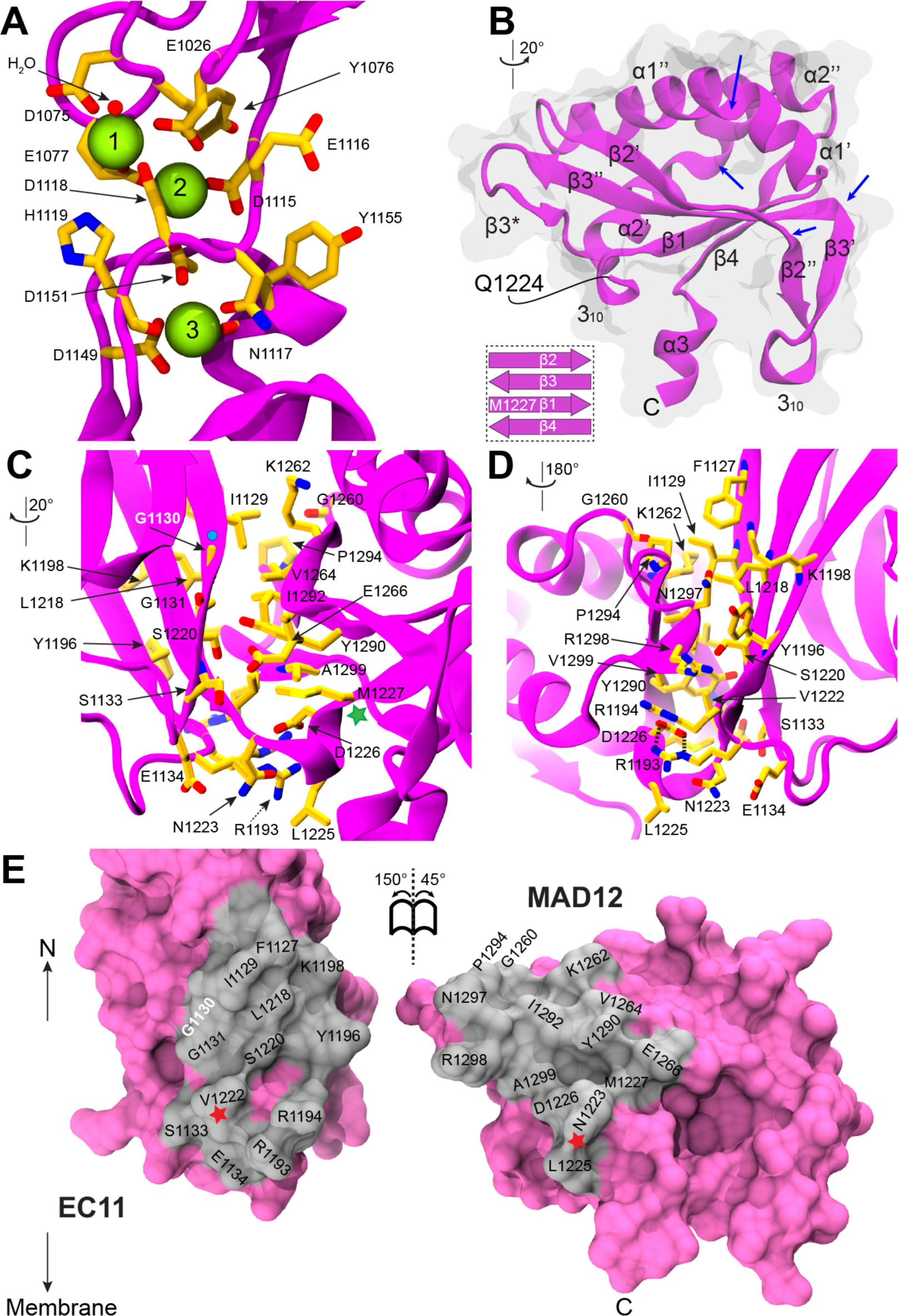
Structural details of EC10-MAD12 fragment. (***A***) Detail of EC10-11 linker region. Ca^2+^ ions are shown in green and Ca^2+^-coordinating side chains in yellow sticks. (***B***) MAD12 in ribbon representation with secondary structure elements labeled. Blue arrows indicate kinks. Inset shows β sheet arrangement. (***C*-*D***) Detailed view of the intramolecular interactions at the EC11-MAD12 interface. Residues are shown in yellow sticks and labeled; some backbone atoms are omitted for clarity. Cyan circle indicates site involved in inherited deafness (p.G1130R). Green star indicates the first residue of MAD12. (***E***) Interaction surface exposed (gray). Red star indicates connection point. Deafness mutation site p.G1130 is highlighted in white. Rotations in panels (*C*) and (*E*) are indicated with respect to view of *ss* PCDH15 EC10-MAD12 in Fig. 1 *B*. Rotation in panel (*D*) is 9 with respect to view in panel (*C*).

The overall architecture of MAD12 is similar to that observed for the ferredoxin-like fold in the Nup54 αβ domain (PDB ID 5C2U, core RMSD 1.64 Å) (95). MAD12 has four β strands and two kinked α helices in a βαββαβ sequence that ends in an α-helical linker (α3) connecting to PCDH15’s transmembrane helix, with kinks dividing α1 into α1’ and α”, as well as,α into α2’ and α2” (Figs. 1 *C* and 2 *B*). Both MAD12 and Nup54 have a short β strand preceding α2’ (labeled β3*) that forms a β-hairpin with the end of β3. There are also key differences between these folds, the most significant being related to kinks in MAD12 that divide β2 into β2’ and β2”, and β3 into β3’ and β3”. The unique β2”-β3’ hairpin, bridged by a 3_10_ helix, bends to form a “hook” reaching in the direction of the membrane, thereby establishing contacts that support the α3 linker connecting to PCDH15’s transmembrane helix (Figs. 1 *B* and 2 *B*, Fig. S1 *B* and *D*).

The L-shaped structure of the PCDH15 EC10-MAD12 protomer harbors various intramolecular contacts between EC11 and MAD12 involving surface residues located at the EC11-MAD12 linker, at the beginning of β2’, at the end of β3”, and at the β3”- β3*-α2’ connection (Fig. 2, *C*-*E*). The interaction surface is ~640 Å^2^ and involves both hydrophilic and hydrophobic contacts. A salt-bridge formed by highly conserved residues p.R1193 (at the E’F’ loop of EC11) and p.D1226 (at the 3_10_ helix in the EC11-MAD12 linker) flanks the external side of the interface’s “elbow”. At the core of the contacts there are several hydrophobic residues that stabilize the interface, including but not limited to p.F1127, p.I1129, p.L1218, and p.V1222 in EC11, as well as p.V1264, p.Y1290, p.I1292, p.P1294, and p.A1299 in MAD12. The hydrophobic nature of these residues is strictly conserved across species (Fig. 2 *C* and Fig. S2), with some variations in sidechain size, which suggests that this intramolecular contact is somewhat robust to residue substitutions as long as their hydrophobicity is maintained. Interestingly, a missense mutation (p.G1130R) that causes inherited deafness (96) alters a site at this interface (Fig. 2, *C*-*E* and Fig. S1 *C*). The sidechain of the mutated residue would not point towards the interface, but would likely destabilize the hydrophobic core of EC11. This suggests that disruption of the structural integrity of EC11 and the intramolecular EC11-MAD12 contacts might be deleterious for PCDH15 function.

### Pig PCDH15 EC10-MAD12 Forms Parallel *cis* Dimers *in crystallo*

The two protomers observed in the asymmetric unit form a parallel *cis* dimer with three contact points (Fig. 3). The first one involves interactions between the two copies of repeat EC10, which face each other bringing together residues on the surface of their o-strands A, G, and F (Fig. 3, *A*, *C*, and *E*). The surface area of this point of contact is small (~185 Å^2^) with interactions mediated by charged and polar residues forming hydrogen bonds and salt-bridges (p.E1017, p.E1018, p.R1020, p.R1085, p.K1108, and p.Y1110). There is strict conservation of charge for all five charged sites in the pig construct, while p.Y1110 is conserved only in mammals, and replaced by hydrophobic residues in birds, reptiles, and some fish (Fig. S2). Details of this contact may minimally vary across species. Interestingly, the dynamics of this EC10-EC10 *cis* interface might be allosterically controlled by Ca^2+^-binding to the EC10-11 linker positioned right below it, thereby providing some Ca^2+^-dependent rigidity at the lower end of the tip link.

**FIGURE 3.**
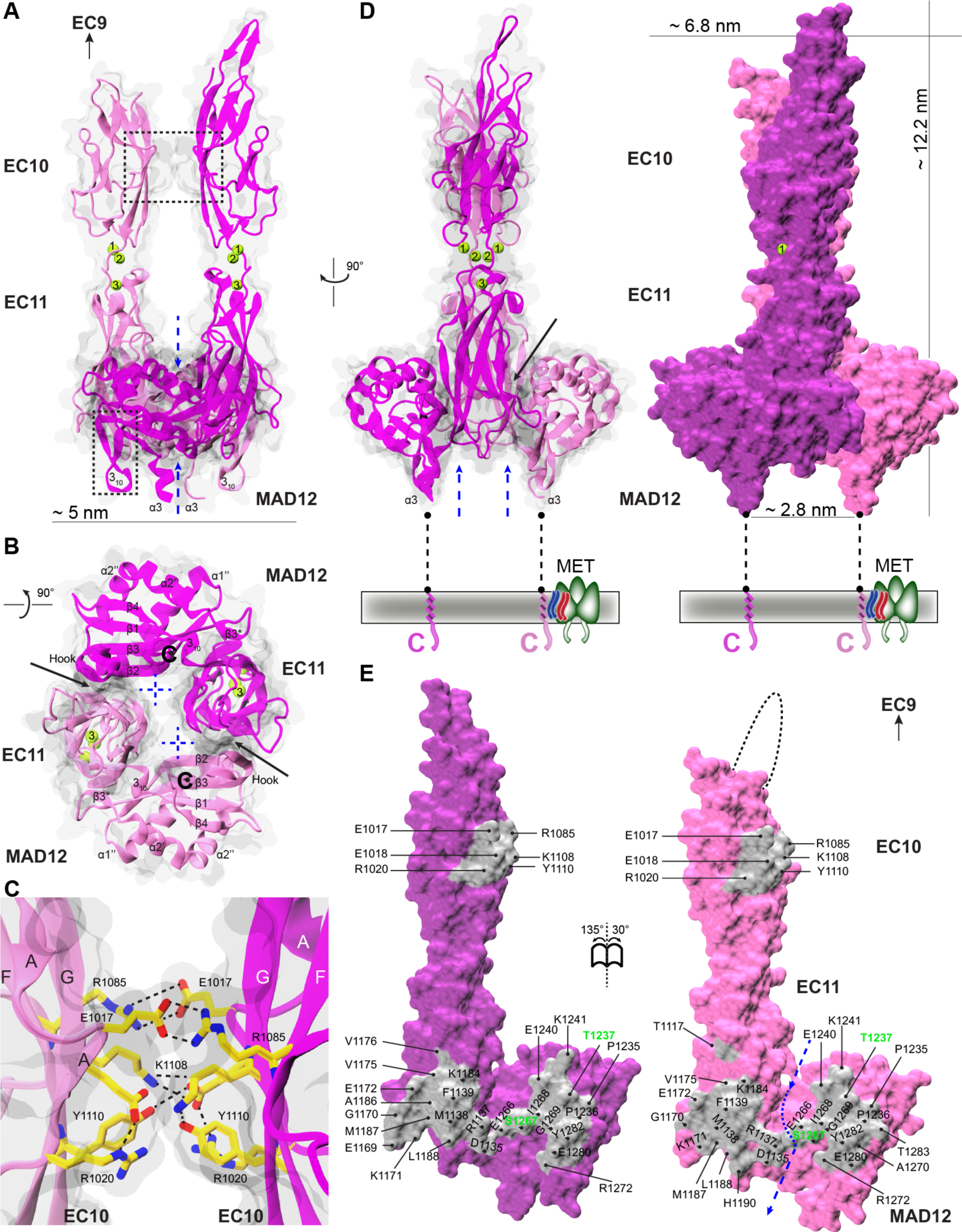
Dimerization interface in *ss* PCDH15 EC10-MAD12. (***A*-*B***) Front and bottom views of *ss* PCDH15 EC10-MAD12 dimer in ribbon (magenta and mauve) with a transparent molecular surface. Black-dashed boxes indicate EC10-EC10 interface (*A*, top) and MAD12’s hook (*A*, bottom). Dashed blue arrows indicate opening crossed by p.R1137. Solid black arrows indicate binding interface between protomers. (***C***) Detail of EC10-EC10 interface (black box in *A*). Protein backbone is shown in ribbons and relevant residues are shown as yellow sticks. (***D***) Side view of dimer in ribbon (left) and opaque surface (right) representations. (***E***) Interaction surfaces exposed and shown in gray with interfacing residues labeled. Missing loop in one protomer is indicated by dashed-black line. Deafness associated sites are labeled in green.

The other two contact points, also recently reported in (49), are equivalent and involve the tail end of one of the L-shaped molecules interacting with the elbow of the other, together forming a closed ring arrangement with some additional contacts stemming from residues pointing towards the center of the ring on the membrane-proximal end of the dimer (Fig. 3, *A*-*E* and Fig. S4). Total surface area for these interfaces is ~975 Å^2^, with hydrophobic and hydrophilic residues from EC11’s A’B’ loop (p.R1137, p.M1138 and p.F1139), C’D’ loop (p.E1172),β-strand D’ (p.V1175), the end of β-strand E’ (p.K1184 and p.A1186) and the beginning of the E’F’ loop (p.L1188), as well as residues from MAD12’s s1’ helix (p.P1236, p.T1237, and p.E1240) and its β-strands β2’ (p.E1266) and β3’ (p.E1280 and p.Y1282). Residue p.T1237, which forms a hydrogen bond with p.E1172 at this interface, is associated to inherited deafness. The mutation p.T1237P is correlated with a deafness phenotype, although causality has not been established (97). Interestingly, the ring-like arrangement (Fig. 3 *B*) features an opening exposed to solvent and crossed by p.R1137 residues from the two protomers. These sidechains interact with either the backbone carbonyl group of p.D1135 or with the sidechain of p.E1266 from the neighboring EC11. These interactions bring AB loops from each EC11 protomer together to stabilize the dimeric interface. Charge is again strictly conserved throughout all the species analyzed (Fig. S2), while hydrophobicity is less conserved for two of the sites (p.1138 and p.1188). The total surface area involved in the potential *cis* dimerization interface, including all three points of contact (~1160 Å^2^), and the sequence conservation of residues involved in it suggests that parallel dimerization might be robust and relevant for PCDH15 function.

### Pig and Mouse PCDH15 EC10-MAD12 Form Dimers in Solution

To determine the oligomeric state of PCDH15 EC10-MAD12 in solution and to test the propensity of this fragment to dimerize, as seen in the X-ray crystal structure, we used pig and mouse protein fragments for analytical ultracentrifugation (AUC) experiments, as well as size-exclusion chromatography (SEC) coupled to multi-angle laser light scattering (MALLS) and small-angle X-ray scattering (SAXS). Sedimentation velocity AUC data for the refolded *ss* PCDH15 EC10-MAD12 (produced in bacteria; theoretical monomeric mass of 39.96 kDa) shows a major peak with a sedimentation coefficient (*S*) of ~ 4, and a minor peak at *S* ~ 3 (Fig. 4 *A*). From these AUC measurements, the molar mass of the major peak is ~60 kDa, yet uncertainty in buffer density, viscosity, specific molar volume, and friction coefficients precluded a more accurate estimate. Nevertheless, we assigned these peaks to dimer and monomer forms, respectively. Refolded *mm* PCDH15 EC10-MAD12 (produced in bacteria; theoretical monomeric mass of 39.97 kDa) also showed similar peaks (Fig. 4 *A*). Consistent with AUC, although less conclusive, SEC-MALLS experiments suggest that the *ss* and *mm* PCDH15 EC10-MAD12 proteins are dimeric in solution (M_w_ = 62.13 kDa ± 0.058% and 61.13 kDa ± 0.043%, respectively) (Fig. S5, *A* and *B*). Molecular weight estimates from SEC-SAXS experiments for both the pig and the mouse fragments (76.20 kDa and 74.92 kDa; 4.7% to 6.3% discrepancy with sequence-derived molecular weight) are in better agreement with the expected mass for dimeric PCDH15 EC10-MAD12 in solution.

**FIGURE 4.**
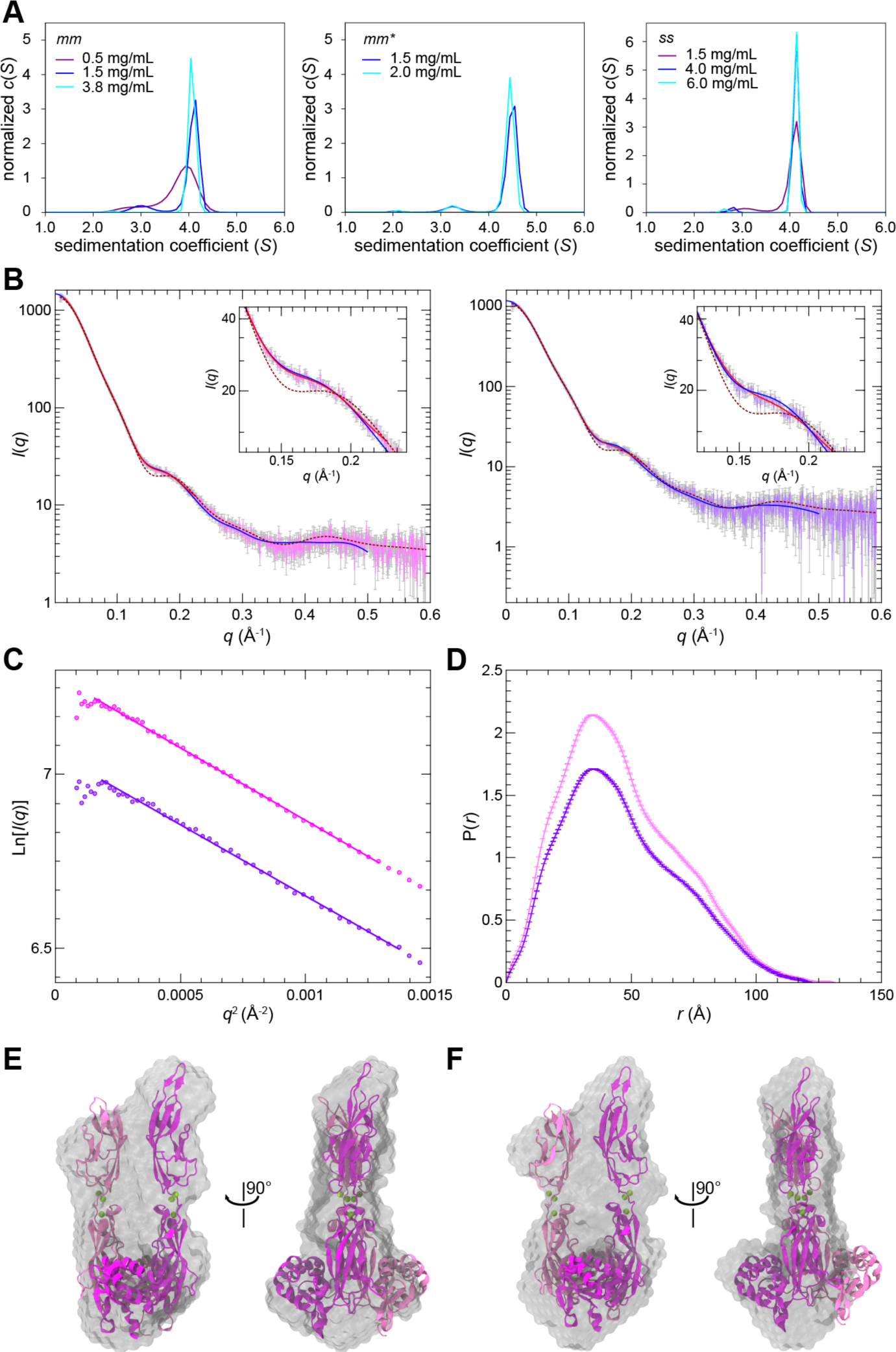
PCDH15 EC10-MAD12 forms a parallel dimer in solution. (***A***) AUC experiments for mouse protein refolded from bacteria (*mm*, left panel), mouse protein purified from mammalian cells (*mm*\*, middle panel), and pig protein refolded from bacteria (*ss*, right panel). Peaks at *S* > 4 represent dimeric states. (***B***) X-ray scattering intensity as a function of the scattering vector *q* (SAXS profile) for pig (magenta, left panel) and mouse (purple, right panel) PCDH15 EC10-MAD12. Predicted scattering intensities from the structure (6BXZ) obtained with FoXS are shown in maroon-dashed lines (χ^2^ = 5.09 / 3.42), while theoretical scattering curves obtained from *ab initio* modeling (DAMMIF) and from flexible refinement with SREFLEX are shown in red (χ^2^ = 1.44 / 1.14) and blue (χ^2^ = 1.64 / 1.23), respectively. (**C**) Guinier plot of the low *q* region of the SAXS data for *ss* and *mm* (magenta and purple circles, respectively) PCDH15 EC10-MAD12. Magenta and purple solid lines show linear fits from which the gradient of the slope (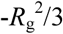) was used to estimate *R*_g_. (***D***) Real-space pair distribution function P(*r*) from SAXS data for *ss* and *mm* PCDH15 EC10-MAD12 (magenta and purple). (***E***-***F***) Superposition of the *ss* PCDH15 EC10-MAD12 structure (6BXZ) with refined low-resolution bead models 13 obtained for pig (44 ± 3 Å) and mouse (37 ± 3 Å) PCDH15 EC10-MAD12 data.

Similarly, glycosylated *mm*\* PCDH15 EC10-MAD12 produced in a mammalian expression system behaves as a dimer in solution. AUC experiments show that *mm*\* PCDH15 EC10-MAD12 is predominantly a dimer with a calculated mass of 89.4 ± 0.3 kDa. The *c*(*S*) distribution (Fig. 4A) also shows a small monomer peak corresponding to a mass of 54.6 ± 2.1 kDa). The expected mass is ~55 kDa, as determined by SDS-gel analysis, with the additional mass due to post-translational modifications on the protein (Fig. S3). Estimates from SEC-MALLS (130.3 kDa ± 0.299%) are consistent with the AUC data and with the expected mass of a dimeric *mm*\* PCDH15 EC10-MAD12 in solution. Together, this data strongly support a dimeric arrangement at the membrane proximal region of PCDH15, as also recently reported for similar protein fragments (48,49).

Data from SAXS experiments can also be used to obtain information about the size and shape of a protein in solution. The radius of gyration (*R*_g_) for both *ss* and *mm* PCDH15 EC10-MAD12 was estimated using two approaches. First, the Guinier analysis was used to obtain *R*_g_ values of 35.37 ± 0.27 Å and 35.53 ± 0.53 Å, respectively (Fig. 4, *B* and *C*; Table S3). In the second approach, the indirect Fourier transform of the SAXS profile yielded *R*_g_ values of 36.30 ± 0.09 Å for the pig and 36.34 ± 0.08 Å for the mouse protein, with maximum dimensions (*D*_max_) of 131 Å and 122 Å, respectively (Fig. 4 *D* and Table S3). These estimates are in excellent agreement with each other and with the *R*_g_ obtained using the *ss* PCDH15 EC10-MAD12 structure (34 Å), suggesting that the overall shape of the dimer observed in the crystallographic structure is maintained in solution. However, comparison of the SAXS data to X-ray intensities modeled from the crystal structure revealed some discrepancies reflected in large χ^2^ values obtained for the fitting (5.09 and 3.42 for *ss* and *mm* PCDH15 EC10-MAD12, respectively) and by a clear dip in the *q*-region 0.1 – 0.25 Å^−1^ observed in the modeled intensities but absent in the experimental data (Fig. 4 *B*). These discrepancies may indicate loss of symmetry in solution and may arise from the relaxation and re-accommodation of the dimer interface and “wobbling” of the protein. Analysis of the Kratky plot (Fig. S5 *C*) indicates that while both proteins are folded in solution, they exhibit significant flexibility, likely arising from inter-repeat motion or the presence of a C-terminal tail including residues p.I1342 to p.E1353 (not seen in the structure) and a His-tag used for protein purification.

The presence of a flexible or conformationally heterogeneous dimer in the samples was further highlighted by *ab initio* modeling of the SAXS data. Thirty models produced for shape reconstruction considering P1 symmetry and prolate anisometry (see Methods) were averaged to generate final models that were refined against the SAXS data for each sample using DAMMIF/DAMMIN/DAMSTART (73,75). Predicted X-ray intensities of the resulting bead models are in good agreement with the experimental data (χ^2^ =1.45 for pig and χ^2^ =1.15 for mouse) and with the crystallographic structure (Fig. 4, *E*-*F*). However, the bead models superposed to the crystallographic structure highlight the lack of symmetry of the dimer in solution and the likely flexibility of the EC10-EC10 interface.

To determine if the crystallographic structure could be re-accommodated in a way that fits the experimental data, we carried out normal-mode analysis on it followed by fitting to the SAXS data with SREFLEX (78) for data from both pig and mouse samples. X-ray intensities calculated from these models fit the experimental data better than the crystal structure (χ^2^ =1.64 for pig and χ^2^ =1.23 for mouse), and lack the dip in the *q*-region 0.1 – 0.25 Å^−1^. The adjusted models have an overall root-mean-square-deviation (RMSD) for Cα atoms of ~7 Å when compared with the crystallographic structure. These models show changes in the orientation and rotation of the ECs with loss of symmetry and with one EC10 repeat tilted and leaning towards the adjacent, straighter EC10 (Fig. S5 *D*). Overall, SAXS data suggest a flexible and perhaps asymmetric arrangement of EC10 repeats while strongly supporting a dimeric PCDH15 EC10-MAD12 in solution.

### Steered Molecular Dynamics Simulations Predict the Mechanics of *ss* PCDH15 EC10-MAD12

The topology and structure of MAD12 has important implications for the transmission of force to the inner-ear transduction channel. Unlike the β-sandwich strand Greek-key fold of EC repeats, in which N- and C-termini are at opposite ends of the folding unit, the N-terminus of MAD12’s βαββαβ fold is spatially nearby its C-terminal end (within ~5 Å). While stretching and mechanical unfolding of an EC repeat requires large forces to shear-out β-strand A or G (98–100), unfolding of MAD12, when pulling from the N- and C-termini, would require unzipping of β4 from β1 and subsequent unraveling of α2. Experiments and simulation have suggested that unzipping of β strands is easier than shearing (101–105), indicating that unfolding of MAD12 may require less force than the force needed to unfold EC repeats.

To explore the elastic response of *ss* PCDH15 EC10-MAD12, its unfolding pathway, and its mechanical strength, we carried out SMD simulations in which force was applied through independent springs attached to the ends of the protein fragment (83–85,106,107). The free ends of the springs were in turn moved at constant speed in opposite directions, mimicking *in vivo* force application. Simulation systems, encompassing up to ~279,000 atoms (Table S4), included explicit solvent and ions along with either a monomer or a dimer of *ss* PCDH15 EC10-MAD12. Stretching of the monomer at 0.1 nm/ns revealed quick unraveling of α3, followed by opening of the EC11-MAD12 interface (separation of p.R1144 from p.E1244) and unrolling of MAD12 accompanied by a 90p swing of MAD12’s hook (Fig. 5 *A* and Fig. S6, *A* and *B*, and Video S1). A final unfolding event involved both unzipping of β4 from 41 with rupture of backbone hydrogen bonds, and splitting of two salt bridges formed during the SMD trajectory (p.D1286-p.R1334 and p.R1271-p.E1332) (Fig. 5, *A* and *B*, Fig. S6, *A*-*G*). Rupture of this “electrostatic lock”, which linked β4 with β2” (hook) and with β3”, correlated with the largest force peak (477.9 pN) monitored throughout the simulation (Fig. S6 *C*). Further unfolding of MAD12’s s2 ensued at drastically smaller forces, without noticeable unfolding of EC10 or EC11. A similar scenario was observed in the slowest stretching simulation at 0.02 nm/ns (S1e, Table S4), with the magnitude of force peaks decreasing at slower stretching speeds as expected (108). Monomer force peaks at all stretching speeds were associated to MAD12 unfolding and were smaller in magnitude than unfolding forces reported for other CDH23 and PCDH15 EC repeats stretched under similar simulation conditions (Fig. 5 *G*) (45,46,100). In addition, MAD12’s unfolding force peaks were smaller than predicted unbinding force peaks for the PCDH15-CDH23 monomeric complex (43,109), suggesting that a single tip-link handshake bond formed by EC1-2 repeats of CDH23 and PCDH15 can withstand similar or larger forces than MAD12.

**FIGURE 5.**
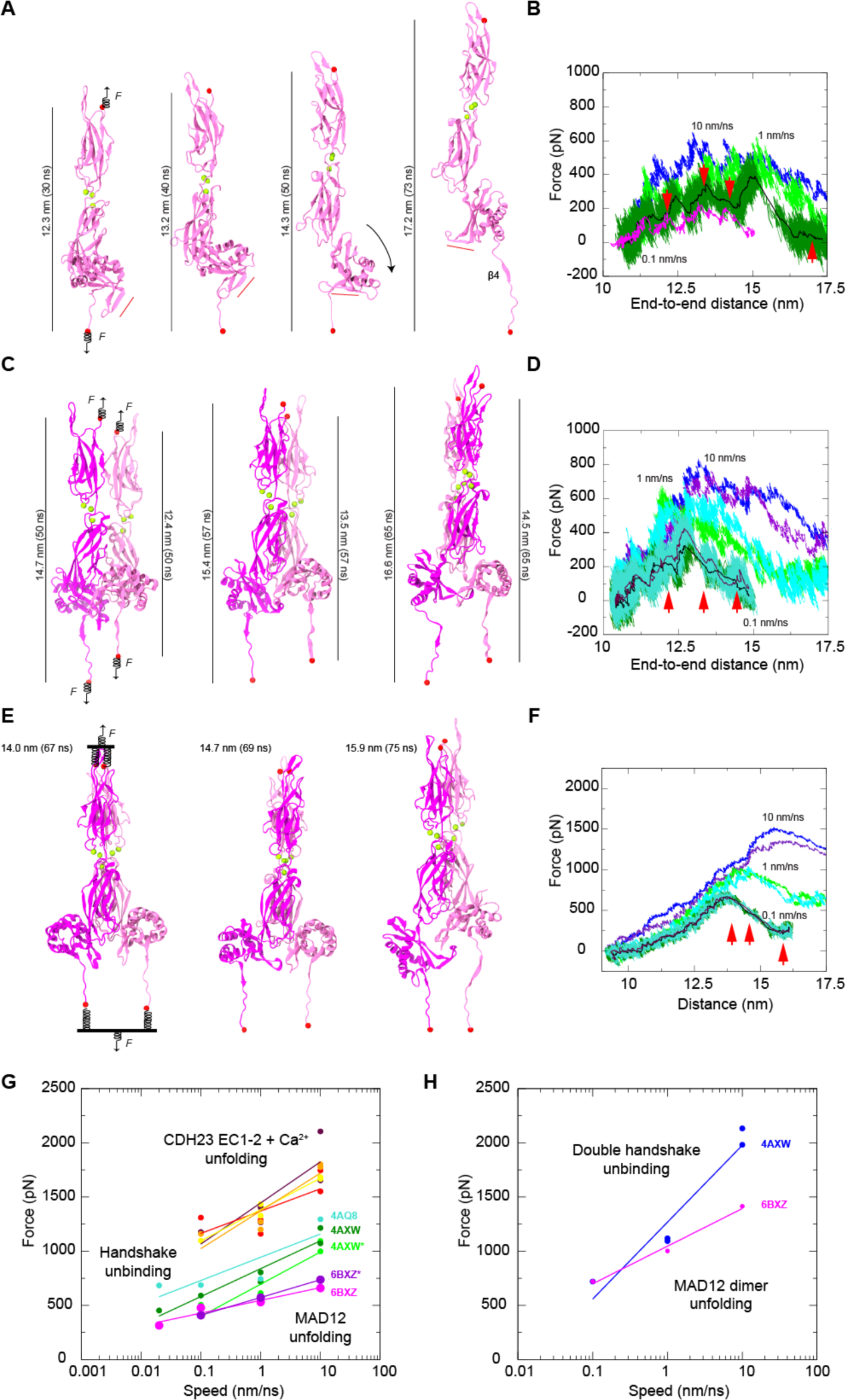
Constant-velocity SMD simulations of *ss* PCDH15 EC10-MAD12. (***A***) Snapshots of monomeric *ss* PCDH15 EC10-MAD12 stretching from simulation S1d (0.1 nm/ns). Protein is shown in mauve ribbon representation, Ca^2+^ ions as green spheres, and stretched terminal C atoms as red spheres. Springs indicate position and direction of applied forces. Red bar illustrates direction of PCDH15’s MAD12 Hook. (***B***) Force applied to C-terminus versus end-to-end distance for constant velocity stretching of monomeric *ss* PCDH15 EC10-MAD12 at 10 nm/ns (S1b, blue), 1 nm/ns (S1c, light green), 0.1 nm/ns (S1d, dark green; 1-ns running average shown in black) and 0.02 nm/ns (S1e, 1-ns running average shown in magenta). Red arrowheads indicate time-points for S1d illustrated in (*A*). (***C***) Snapshots of dimeric *ss* PCDH15 EC10-MAD12 stretching from simulation S2d as in (*A*), with subunits in magenta and mauve. (***D***) Force applied to N- and C-termini of one subunit (mauve in *C*) versus end-to-end distance for constant-velocity stretching of dimeric *ss* PCDH15 EC10-MAD12 at 10 nm/ns (S2b, blue and indigo), 1 nm/ns (S2c, light green and cyan), and 0.1 nm/ns (S2d, dark green and turquoise; 1-ns running average shown in black and maroon). Red arrowheads indicate time-points illustrated in (*C*). (***E***) Snapshots of dimeric *ss* PCDH15 EC10-MAD12 stretching from simulation S2g as in (*C*). Stretching was carried out by attaching two slabs to springs that were in turn attached to the terminal ends of each protein. Slabs were moved in opposite directions through individual springs. (***F***) Force applied to each of the slabs versus slab separation for constant-velocity stretching of dimeric *ss* PCDH15 EC10-MAD12 at 10 nm/ns (S2e, blue and indigo), 1 nm/ns (S2f, light green and cyan), and 0.1 nm/ns (S2g, dark green and turquoise; 1-ns running average shown in black and maroon). (***G***) *In silico* force peak maxima versus stretching speed for CDH23 EC1-2 unfolding (red, maroon, yellow, and orange) (100), for CDH23 EC1-2 and PCDH15 EC1-2 handshake unbinding (cyan-4AQ8, dark green-4AXW, and light green-4AXW* [after 1 µs-long equilibration]) (43,109), and for MAD12 unfolding (magenta-6BXZ for simulations S1b-d; purple-6BXZ* for averages obtained for simulations S2b-d). (***H***) *In silico* force peak maxima versus stretching speed for unbinding of pairs of CDH23 EC1-2 and PCDH15 EC1-2 handshakes (blue) (43) and for unfolding of MAD12 using slabs (magenta, simulations S2e-g).

Stretching of the *ss* PCDH15 EC10-MAD12 dimer was carried out using two constant-velocity SMD protocols and revealed a more complex response. First, we applied forces in opposite directions through independent springs attached to each of the four ends of the protein fragments (Fig. 5 *C*). At the slowest stretching speed (0.1 nm/ns), one of the subunits’ β4 detached from β1 as MAD12 was separating out from EC11 and unrolling, which correlated with a broad and rather undefined force peak (not shown). In the other subunit MAD12 separated from EC11 and unrolled after unzipping β4 from β1 and rupture of the electrostatic lock described above, which correlated with a clear force peak (Fig. 5 *D*). In a second approach, pairs of N- or C-termini were connected to virtual slabs through independent springs (Fig. 5 *E*). The slabs were in turn connected through springs to SMD atoms moving at constant speeds in opposite directions. In the simulation that used this approach at the slowest stretching speed (0.1 nm/ns), MAD12 from one subunit separated from EC11 and unrolled before β4 detached from β1, while MAD12 from the other subunit remained partially attached to EC11, with unzipping of β4 from β1 happening first (Fig. 5 *E* and Fig. S7, Video S2). In all cases, additional intermolecular interactions formed throughout trajectories and MAD12’s hook swung as monitored during stretching of the monomer of *ss* PCDH15 EC10-MAD12. Average unfolding force peaks of MAD12 were smaller than unfolding force peaks of PCDH15 EC repeats and unbinding force peaks of the PCDH15 EC1-2-CDH23 EC1-2 bond, again suggesting that MAD12 is mechanically weak, with the tip-link bond being of similar or larger strength at the stretching speeds tested in this study (Fig. 5, *G* and *H*).

### Secondary Structure Analyses and Modeling Predict MAD Folds in Other Cadherins

The topology and structure of MAD12 has important implications for other cadherins as well. The high structural similarity observed between MAD12 and Nup54 (95), despite their very poor sequence identity (Figs. S8 and S9), led us to search for other βαββαβ domains that might be hidden within the cadherin superfamily (Fig. 6). We looked for extracellular segments that lacked predicted domain architecture, referred to as unknown regions according to SMART (110). In parallel, we predicted the secondary structure of sequences from all cadherins in the human genome using RaptorX (111), which had correctly predicted the secondary structure of MAD12 and identified Nup54 (PDB 5C2U) (95) as a good template for “homology” modeling. We then focused on sequences predicted to have the βαββαβ fold, and found the pattern in CDH23, PCDH24 (also known as CDHR2), all FAT cadherins, and all CELSRs (Figs. S8 and S10). These predicted βαββαβ-fold containing segments were classified as unknown regions by SMART.

**FIGURE 6.**
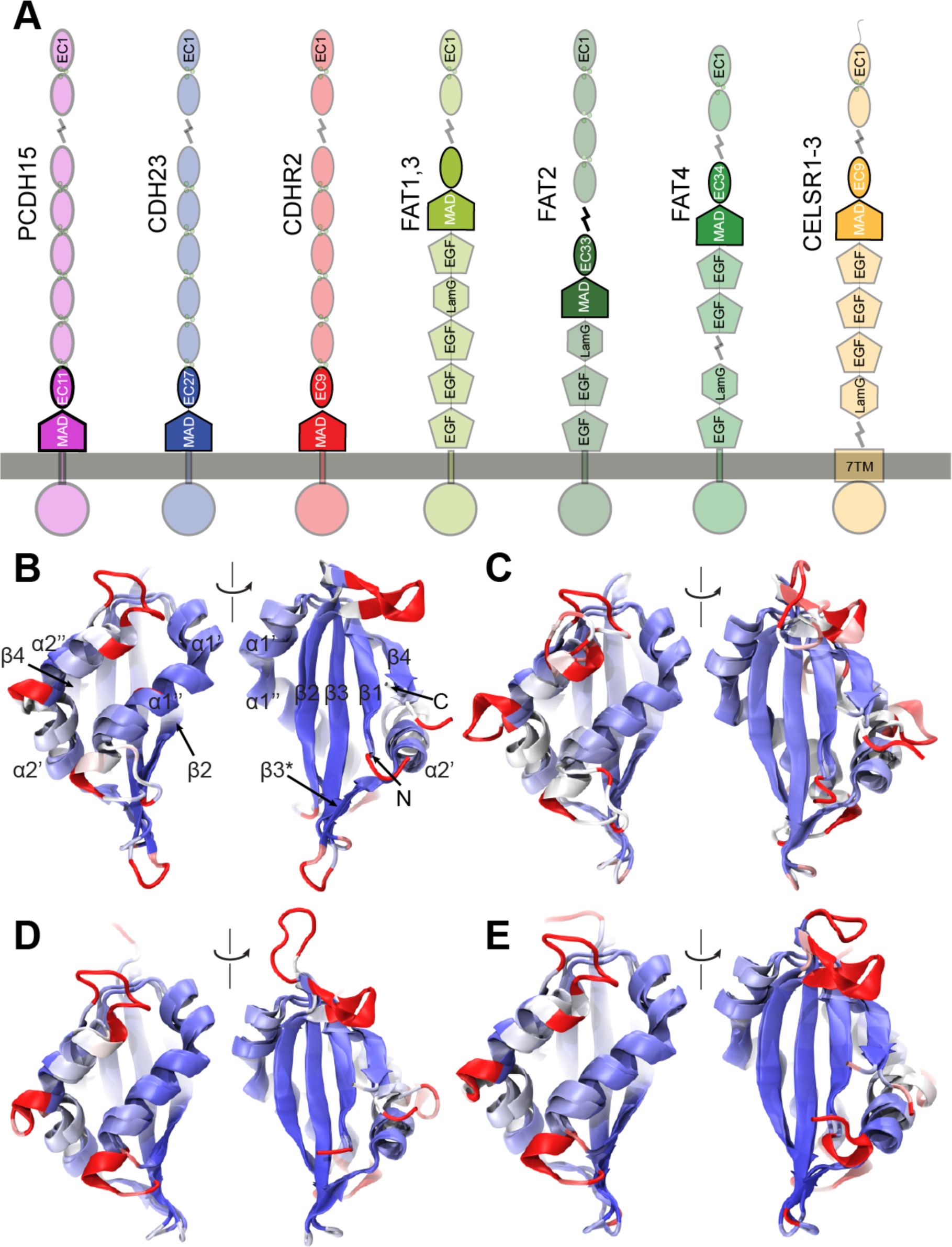
Presence of putative MADs in the cadherin superfamily. (***A***) Domain organization of human cadherins predicted to feature domains similar to PCDH15 MAD12. (***B***) Structural comparison between *xl* Nup54 αβ domain (5C2U) (95) and *ss* PCDH15 MAD12 (PDB ID 6BXZ; front and back views). Molecules are colored according to structural homology per residue (*Q*_res_) (114). Blue color implies high structural conservation, whereas red implies poor structural conservations. Beginning of β1 and α3 were truncated out for better structural alignment. (***C***-***E***) Structural models of MADs from atypical cadherins (*hs* PCDH15 MAD12; *hs* CDH23 MAD28; *hs* PCDH24 MAD10) superposed on the crystal structure of *ss* PCDH15 MAD12 (6BXZ), shown as in *B*.

The potential presence of MADs in CDH23 and PCDH24 is intriguing (MAD28 and MAD10, respectively) (Fig. 6). In both cases, these domains would be located close to their membrane insertion points, linking cadherin EC repeats to their associated transmembrane helices. Both proteins had been identified as members of the Cr-2 subfamily based on sequence similarity of their extracellular tips (1,4,100,112) and the presence of a ferredoxin-fold domain in both these proteins further confirms their structural relationship. Perhaps even more intriguing is the existence of potential MAD folds in human FAT and CELSR cadherins. Analysis of the *drosophila melanogaster* (*dm*) FAT also shows the potential existence of a similar βαββαβ fold (MAD35) (Fig. S10). In all these cases, the predicted “unknown region” is between the last, C-terminal EC and subsequent EGF or LAMG domains (MAD35 for FAT1,4; MAD34 for FAT2; MAD33 for FAT3; and MAD10 for CELSR1-3) (Fig. 6 *A*).

To gain further insights into the structural similarities among cadherin MADs we used RaptorX to create structural models for each of these domains in the human “MAD bearing” cadherins (PCDH15, CDH23, PCDH24, FAT1-4, and CELSR1-3) and the fly sequences (*dm* FAT) mentioned above (11 models) (Fig. 6, *B*-*E* and Fig. S10, *A*-*G*). Visual inspection confirmed the βαββαβ fold for eight of these models (*hs* PCDH15, *hs* CDH23, *hs* PCDH24, *hs* FAT1, *hs* FAT2, *hs* CELSR1, *hs* CELSR2, and *dm* FAT). In addition, we classified models according to the hydrophobicity of their core. Hydrophobic cores without polar or charged residues pointing into it were observed for PCDH15 MAD12, CDH23 MAD28, FAT1 MAD35, and CELSR1-2 MAD10, which also had their charged residues distributed on their surfaces and loops, as expected. Models for PCDH24 MAD10, FAT2 MAD34, and *dm* MAD35 were satisfactory, with one or two polar residues lying at the interface between their hydrophobic cores and loops, which could be easily accommodated by rotamer rearrangement. A structural alignment of these models to *ss* PCDH15 MAD12 using STAMP (113,114) revealed high structural conservation at the core secondary structure elements, and poor structural conservation of loops for the eight models mentioned above (Fig. 6).

The dimeric structure of *ss* PCDH15 MAD12 suggests that other MADs could form homodimers as well. However, part of the dimer interface involves the preceding EC repeat and the β2”–3_10_–β3’ hooks that are not well predicted by RaptorX (see “control” prediction in Fig. 6 *B*). Nevertheless, the β2”–3_10_–β3’ hook observed in *ss* PCDH15 MAD12 might be present in CDH23 MAD28. Although the hook is not observed in our model, CDH23 MAD28 has similar length and distribution of charged residues in this region (Fig. S8). Similarly, in the secondary structure prediction by RaptorX (Fig. S8), the short β3* strand interacting with the preceding EC repeat exist in CDH23 and CELSR1 and 2, thus suggesting an architecture that might be similar to the L-shaped PCDH15 MAD12 favoring dimerization.

To determine whether parallel dimerization of MADs in other cadherins is possible *in silico*, we built models of *hs* CDH23 EC26-MAD28 and *hs* PCDH24 EC8-MAD10 and computed their interaction energies. Homology models for EC repeats were coupled to RaptorX models for MADs using the *ss* PCDH15 EC10-MAD12 structure as a guide. The resulting, models with L-shaped protomers were minimized as dimers and used for “blind” molecular docking to predict binding poses for CDH23 and PCDH24, which resulted in *cis* parallel dimers that are similar to *ss* PCDH15 EC10-MAD12 (Fig. S11). In both cases, the L-shaped architecture of *hs* CDH23 EC26-MAD28 and *hs* PCDH24 EC8-MAD10 favors the formation of a closed ring-like arrangement involving several hydrophobic residues that could stabilize their dimeric interfaces. In addition, free energy binding calculations (via Molecular Mechanics/Generalized Born Surface Area [MM/GBSA]) predict strong binding for these admittedly biased dimers (Table S5), suggesting that MAD-induced parallel dimerization is possible in CDH23 and PCDH24. Taken together, our modeling analyses suggest that MADs are present in all the aforementioned proteins, with some variations in fold at loops and at the β2”–3_10_–β3’ and β3”–β3* connections that might determine their oligomerization states and function.

## CONCLUSIONS

The structural, biochemical, and computational results presented here shed light on three aspects of PCDH15 function. First, the unique architecture and fold of PCDH15’s MAD12, along with SMD simulations of its elastic response, suggest that this domain is mechanically weak, which has implications on how PCDH15 and the tip link may respond to large mechanical stimuli, e.g., loud sound. Second, MAD12 induces parallel dimerization at the membrane proximal end of PCDH15’s extracellular domain, thus establishing constraints on models of mechanotransduction and how the tip link may interact with other parts of the transduction machinery. Last, simulations suggest mechanisms by which force-induced deformation of MAD12 can trigger conformational changes in associated membrane proteins, thus providing testable hypotheses on how PCDH15 may gate the inner-ear transduction channel, should they be directly coupled to each other.

Determining the structure and mechanical properties of PCDH15 is a necessary step to understand its function in the context of inner-ear mechanotransduction (18,40,42,115–120). The βαββαβ (α/β) fold adopted by PCDH15’s MAD12, revealed here and in (49), is distinct and unique when compared to those of all other EC repeats in PCDH15, which are all β-sandwich modules (all-β). Previous single-molecule force spectroscopy experiments and simulations have suggested the existence of a hierarchy of mechanical strengths that depends on both the secondary structure elements of a protein module and the geometry of stretching (101–105,121). All-β proteins in which mechanical unfolding necessitates shearing of β strands are often strong, followed by all-β proteins in which unzipping of β strands is mechanically easier. In contrast, α/β proteins are mechanically weaker than all-β proteins, and all-α modules seem to be the weakest of all. Consistent with this hierarchy, our SMD simulations of the *ss* PCDH15 EC10-MAD12 fragment show that MAD12 unrolls and separates from EC11 at low force, with ensuing unfolding at moderate forces facilitated by the unzipping of MAD12’s β4 from β1 and rupture of an electrostatic lock formed by conserved charged residues at β4, β2”, and β3”. Peak unfolding forces for the α/β MAD12 at all stretching speeds tested here were smaller than those reported for unfolding of various all-β CDH23 and PCDH15 EC repeats with bound calcium (45,46,100). At the low calcium concentrations of the cochlea (122,123), MAD12 may still stochastically unfold before other EC repeats when tip links are exposed to extreme mechanical stimuli.

Intriguingly, our simulations predict that MAD12 unfolds at similar or smaller forces than those required to break a single PCDH15-CDH23 handshake bond *in silico* (43), suggesting that under some conditions, tip links may lengthen significantly due to unfolding without PCDH15-CDH23 unbinding. A stretch of ~120 unfolded residues could easily account for >40 nm lengthening, but we do not know whether MAD12 would refold back into its ferredoxin-like fold in time for the next mechanical stimulus. While it is unclear whether unfolding could occur before unbinding at significantly slower stretching speeds (equivalent to low-frequency, normal to faint sound) than those used in our simulations (close to high-frequency, loud sound), it provides a plausible explanation to observations of long tip links that might have been artificially stretched during staining or freezing for electron-microscopy imaging. Unfolding also provides a more compelling explanation to electrophysiological experiments in which tip links are extended over 100 nm without compromising transduction (124). It is possible that both unfolding of MAD12 and unbinding of PCDH15 from CDH23 are used to provide distinct safety mechanisms that prevent rupture of other transduction machinery components and that are perhaps adapted to different sets of extreme stimuli.

Parallel dimerization mediated by MAD12 presents additional constraints on how PCDH15 stretches and how it communicates force to its transmembrane protein partners. Predicted unfolding forces of MAD12 in the context of the dimer were not significantly different than those predicted for the monomer, and were smaller than those required to break a double handshake stretched *in silico*, suggesting that unfolding can occur before unbinding. It remains to be determined whether a heterotetrameric handshake bond in which parallel contacts at PCDH15 EC2-3 (48) and between the CDH23 chains may strengthen the handshake even further, thus favoring unfolding over unbinding at all stretching speeds in heterotetrameric configurations. Intriguingly, deformation and unfolding of MAD12 during stretching simulations of the dimer were in some cases asymmetric. Whether unrolling and separation of MAD12 from EC11 occur before or after unzipping of MAD12’s β4 from β1 in an asymmetric fashion might be controlled by dimerization, direction of the applied stretching force, and binding partners.

Asymmetry in parallel dimerization of PCDH15 observed in our SAXS data is consistent with raw data from negative staining and electron cryo-microscopy (48,49). While averaged structural models may not fully reflect asymmetry and flexibility, SAXS data suggest that PCDH15 exhibits asymmetry and that in some cases each of the subunits in the dimer may adopt its own conformation. In the absence of force, the likely flexibility of the EC10-EC10 interface may indicate that the conformations at the lower end of PCDH15 are allosterically controlled and stabilized by MAD12 and by the EC10-EC11 Ca^2+^-binding linker, and not necessarily by EC9-EC9 or EC10-EC10 contacts between both protomers (49). MD simulations also suggest asymmetric unrolling in the MAD12-induced dimer upon force application, indicating that in some cases force might be conveyed by each protomer independently and in slightly different ways to their respective binding partners.

Interestingly, PCDH15 seems to interact promiscuously with multiple transmembrane proteins, including TMCs, various TRP channels, and TMHS (125–128). The most compelling data on these interactions comes from the recent structural and biophysical characterization of the PCDH15 MAD12-TMHS complex (49). While transmembrane domains mediate most of the interaction between TMHS and PCDH15, there are contacts between the extracellular loops of TMHS and the α3 helix and β2”– 3_10_–β3’ hook of PCDH15’s MAD12, as well as with the E’F’ loop of PCDH15 EC11. These interactions suggest that TMHS may stabilize the base of MAD12, perhaps preventing unraveling of PCDH15’s s3 helix upon force application and facilitating membrane tensioning by tip links, which in turn could result in gating of nearby transduction channels (129–131). Alternatively, it is tempting to speculate that unrolling of MAD12 and swinging of its hook, as observed in our simulations upon force application, could induce conformational changes, either in TMHS or directly in TMC1, that result in “uncorking” of the inner-ear transduction channel. Further structural studies and SMD simulations of the entire complex will shed light on the molecular details of this process.

Our results have implications beyond PCDH15 and inner-ear mechanotransduction, as we predict the presence of MADS in other large cadherins such as CDH23, PCDH24, FATs, and CELSRs. Both CDH23 and PCDH24 are expected to be tensioned under physiological conditions, as CDH23 is part of the inner-ear tip link and PCDH24 forms brush-border intermicrovillar links (50). Possible MAD-induced dimerization may play a critical role in CDH23 and PCDH24 function. Interestingly, PCDH15 and CDH23 have been recently implicated in brain circuit assembly (132), while FATs and CELSRs are known to be involved in regulation of cell polarity and migration. Perhaps all “MAD-bearing” cadherins play a mechanical role, either in sensory perception, epithelial morphogenesis and function, or in brain development and wiring (3,133).

## SUPPORTING MATERIAL

Eleven figures, four tables, and two videos are available at: xxxxxxx

## AUTHOR CONTRIBUTIONS

P.D. did protein expression, purification, crystallization, and X-ray data collection of*ss* PCDH15 EC10-MAD12. P.D. and M.S. solved the X-ray crystal structure. R.A.S. did cloning, protein expression, purification, and initial crystallization attempts of *ss* PCDH15 EC10-MAD12, which generated crystals for seed stocks. P.D. and D.C. prepared the seed stock for crystallization. P.D. and D.C. carried out protein expression and purification of *ss* and *mm* PCDH15 EC10-MAD12 for SEC-SAXS experiments. R.A.S. analyzed the SAXS data and generated SAXS models. P.D and D.C. did protein expression and purification of *ss* and *mm* PCDH15 EC10-MAD12 for SEC-MALLS and AUC experiments. D.C. carried out SEC-MALLS and AUC experiments and did data analysis. Y.N. did cloning, expression, and purification of *mm*\* PCDH15 EC10-MAD12 in mammalian cells, carried out AUC and SEC-MALLS with this sample, and did data analysis. P.D. and D.C. did structural and sequence analyses of MADs and Nup54. D.C. did bioinformatics analyses and generated models of MADs. P.D. built homology models of *hs* CDH23 EC26-MAD28 and *hs* PCDH24 EC8-MAD10 and did molecular docking and MM/GBSA calculations. M.S. did MD simulations. M.S. and P.D. wrote and edited the manuscript with help from all co-authors. M.S. trained and supervised all co-authors and assisted in crystal fishing/cryo-cooling, structure refinement, deposition, and data analysis presented in this work.

## ACKNOWLEDGEMENTS

We thank Michal Hammel and Daniel Rosenberg for data collection and initial SEC-SAXS data analysis, Brandon L. Neel for assistance with sequence alignments, and members of the Sotomayor research group for their feedback, training, and discussions. This work was supported by the Ohio State University and the National Institutes of Health - National Institute on Deafness and Other Communication Disorders (NIH/NIDCD R01 DC015271) and by the National Science Foundation through XSEDE (XRAC MCB140226). Simulations were performed at the PSC-Bridges and OSC-Owens (PAS1037) supercomputers. Use of the APS NE-CAT beamlines was supported by NIH (P41 GM103403 & S10 RR029205) and the Department of Energy (DE-AC02-06CH11357) through grants GUP 49774 and 59251. SAXS data collection at the Advanced Light Source SIBYLS beamline is funded through: DOE BER Integrated Diffraction Analysis Technologies program and NIH grants NIGMS P30 GM124169 ALS-ENABLE and S10OD018483. PD is a Pelotonia Fellow and RAS was a Pelotonia fellow.

## SUPPLEMENTAL INFORMATION

**Video S1.** Forced unrolling and unfolding of monomeric *ss* PCDH15 EC10-MAD12. Protein is depicted in cartoon representation, molecular surface is transparent, and calcium ions are shown as green spheres. Water molecules and other atoms are not shown for visualization purposes. The protein is stretched from both ends at 0.1 nm ns^−1^ (simulation S1d in Table S4).

**Video S2.**Forced unrolling and unfolding of dimeric *ss* PCDH15 EC10-MAD12. Protein is depicted as in Video S1. N- and C-terminal ends of each protomer are connected to slabs that are moved in opposite directions at 0.1 nm ns^−1^ (simulation S2g in Table S4).

**FIGURE S1.**
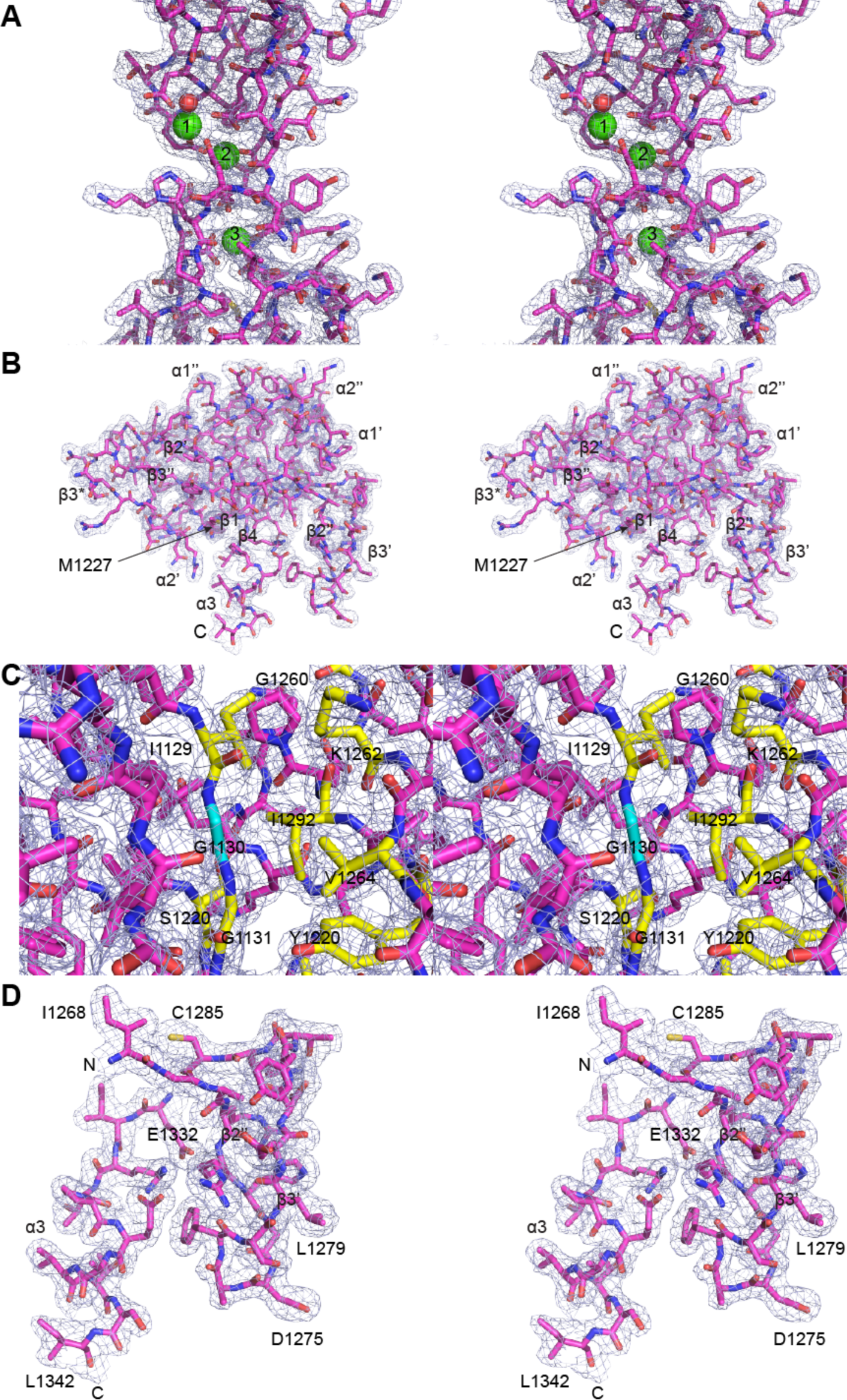
Electron density (light blue) of *ss* PCDH15 EC10-MAD12 regions. (***A***) The EC10-EC11 Ca^2+^- binding linker region (PDB ID: 6BXZ) shown at 0.7 σ. Red sphere represents a water molecule coordinating Ca^2+^ at site 1. (***B***) MAD12 electron density shown at 0.7 σ. (***C***) The EC11-MAD12 protein interface shown at 1.7 σ. Site of deafness-causing mutation (p.G1130R) is in cyan. Some surrounding residues involved in EC11-MAD12 interactions are colored in yellow sticks. (***D***) Electron density for MAD12’s hook and α3 helix shown at 0.7 σ. Residues from β2”, β3’, and β4 are not shown for clarity.

**FIGURE S2.**
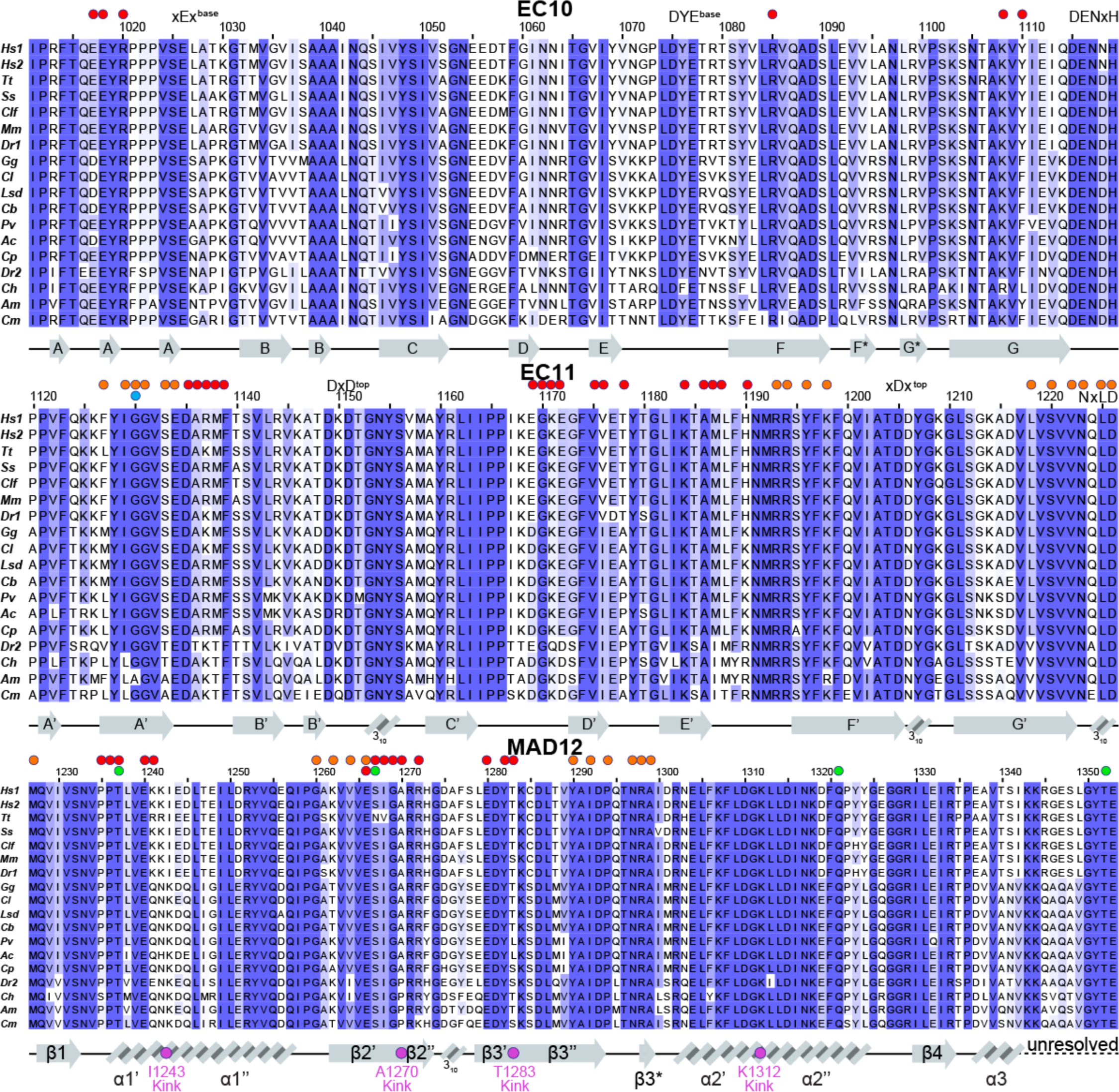
Multiple sequence alignments comparing sequences between EC10, EC11, and MAD12 from 17 different species. Each alignment is colored by % of sequence similarity (see Methods). White represents the lowest similarity, while blue represents the highest. Some columns may not be colored if changes to residue type are present (e.g. polar amino acid to hydrophobic). Secondary structure elements are displayed at the bottom of each alignment. Residues located at the dimer interface and at the EC11-MAD12 interface (determined by PISA) are highlighted by red and orange circles, respectively. Purple circles represent residues at kinks. Cyan and green circles highlight sites involved in hereditary deafness (causal and correlated mutations, respectively). Sequence names for different species are as in Table S1 and were chosen based on sequence availability and taxonomical diversity.

**FIGURE S3.**
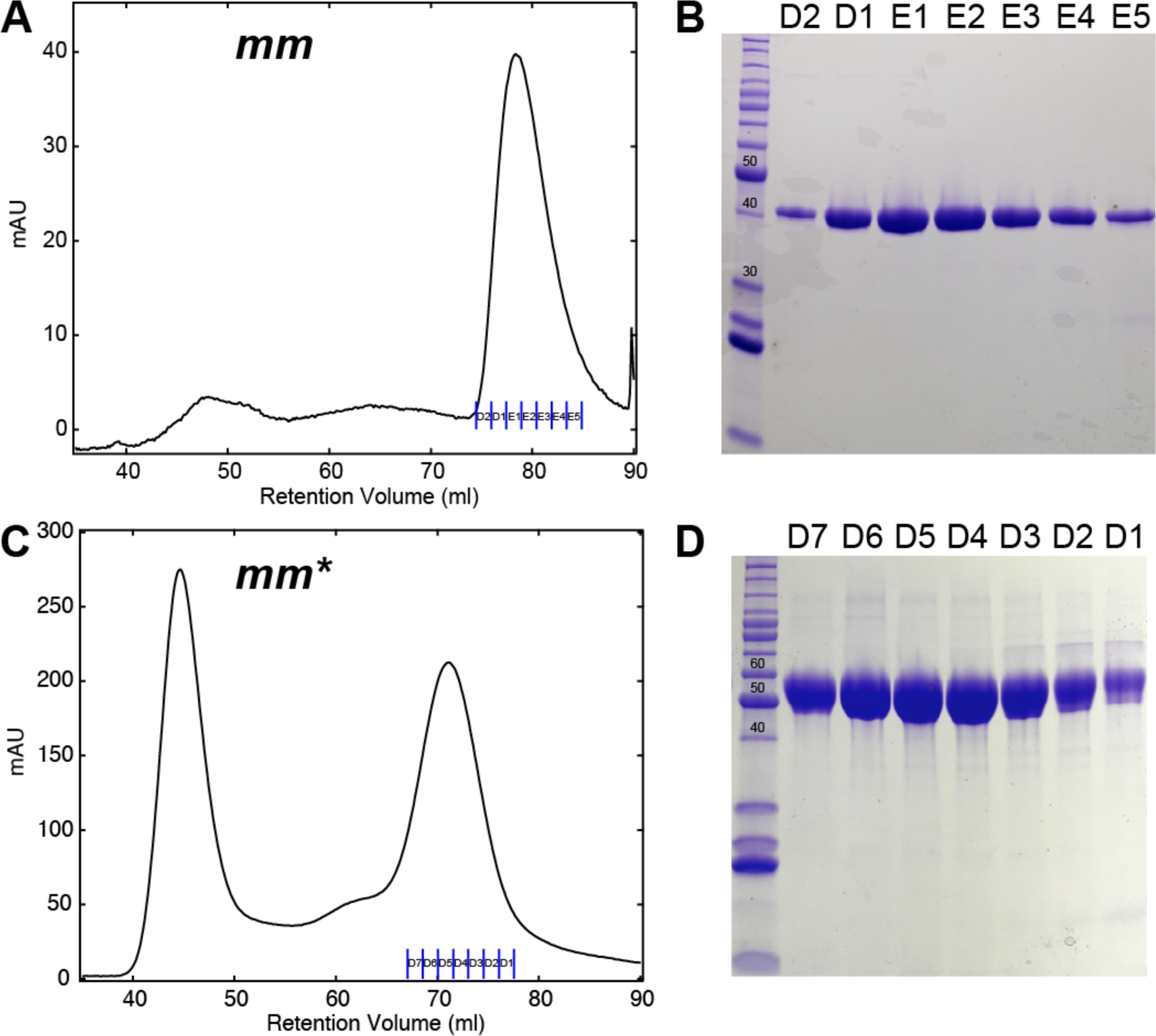
Size exclusion chromatography and SDS-PAGE of *mm* and *mm\** PCDH15 EC10-MAD12. (***A***) Refolded *mm* PCDH15 EC10-MAD12 revealed one main peak at 78.9 ml when purified on a Superdex S200 16/600 column. (***B***) SDS-PAGE analysis of the fractions in the main peak using non-reducing loading dye. (***C***) *mm*\* PCDH15 EC10-MAD12 purified from mammalian cells showed one peak at 44.9 ml (due to aggregation and misfolded protein) and a second, well-behaved peak at 71.2 ml on a Superdex S200 16/600 column. (***D***) SDS-PAGE analysis of the fractions labeled in (*C*) using reducing loading dye. The experimentally determined molecular weight is ~55 kDa.

**FIGURE S4.**
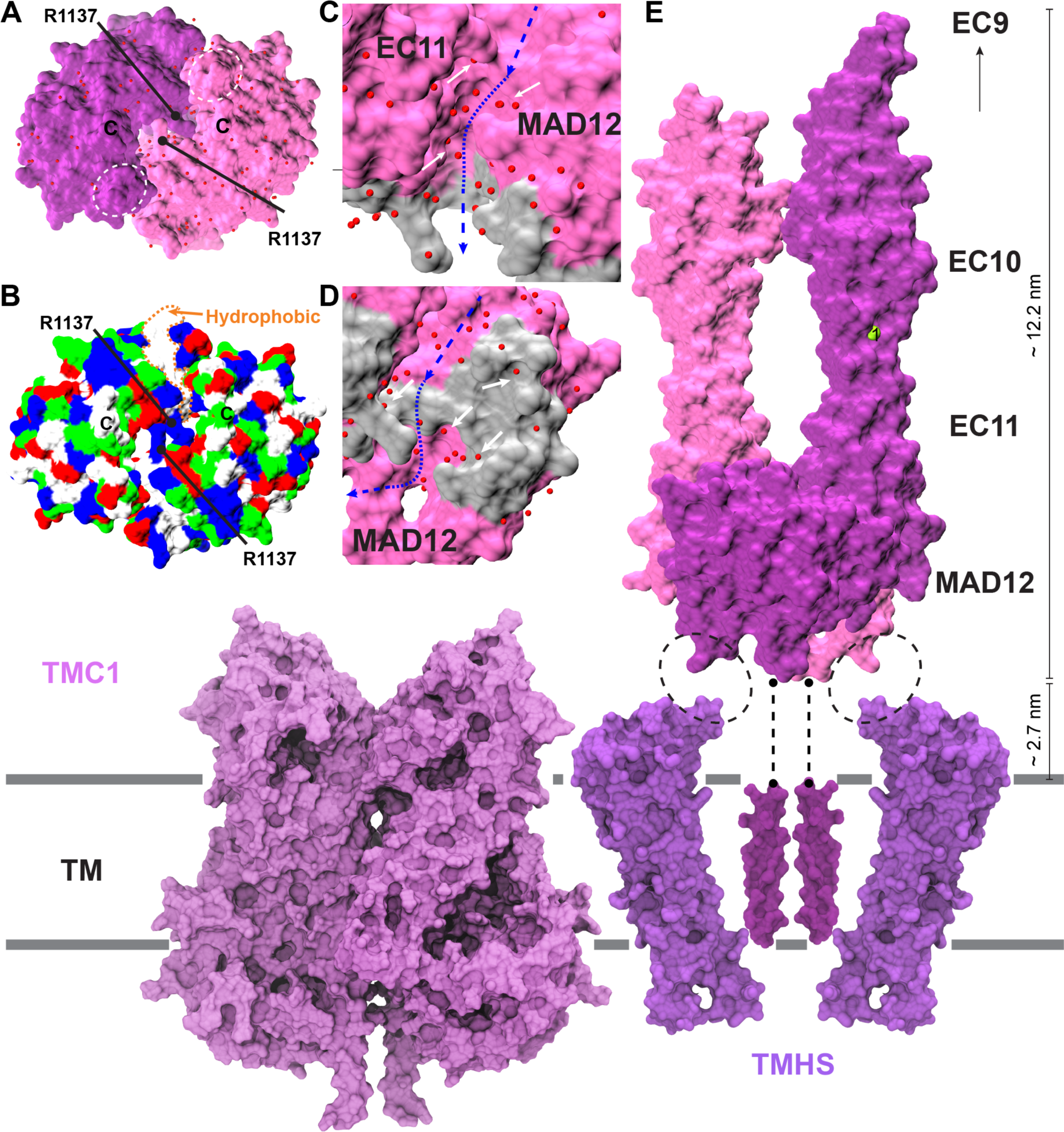
Details of *ss* PCDH15 EC10-MAD12 dimer and schematic representation of the inner-ear mechanotransduction complex. (***A***) Bottom view EC11-MAD12 dimer. White-dashed circles highlight MAD12’s hooks. Water molecules are represented by red spheres. (***B***) Same view as in panel (*A*) colored by residue type (white: hydrophobic; blue and red: charged; and polar residues: green). (***C*-*D***) Top and front views of MAD12’s opening. Interaction surface (gray) as in Fig. 3 *E*. White arrows indicate water molecules located in pockets of the EC11-MAD12 protomer and at the dimer interfaces. (***E***) Schematic representation (to scale) of *ss* PCDH15 EC10-MAD12 dimer, TMHS (purple), and a TMEM-16 based TMC1 model (ligh purple) (134,135). Specific intermolecular interactions are unknown.

**FIGURE S5.**
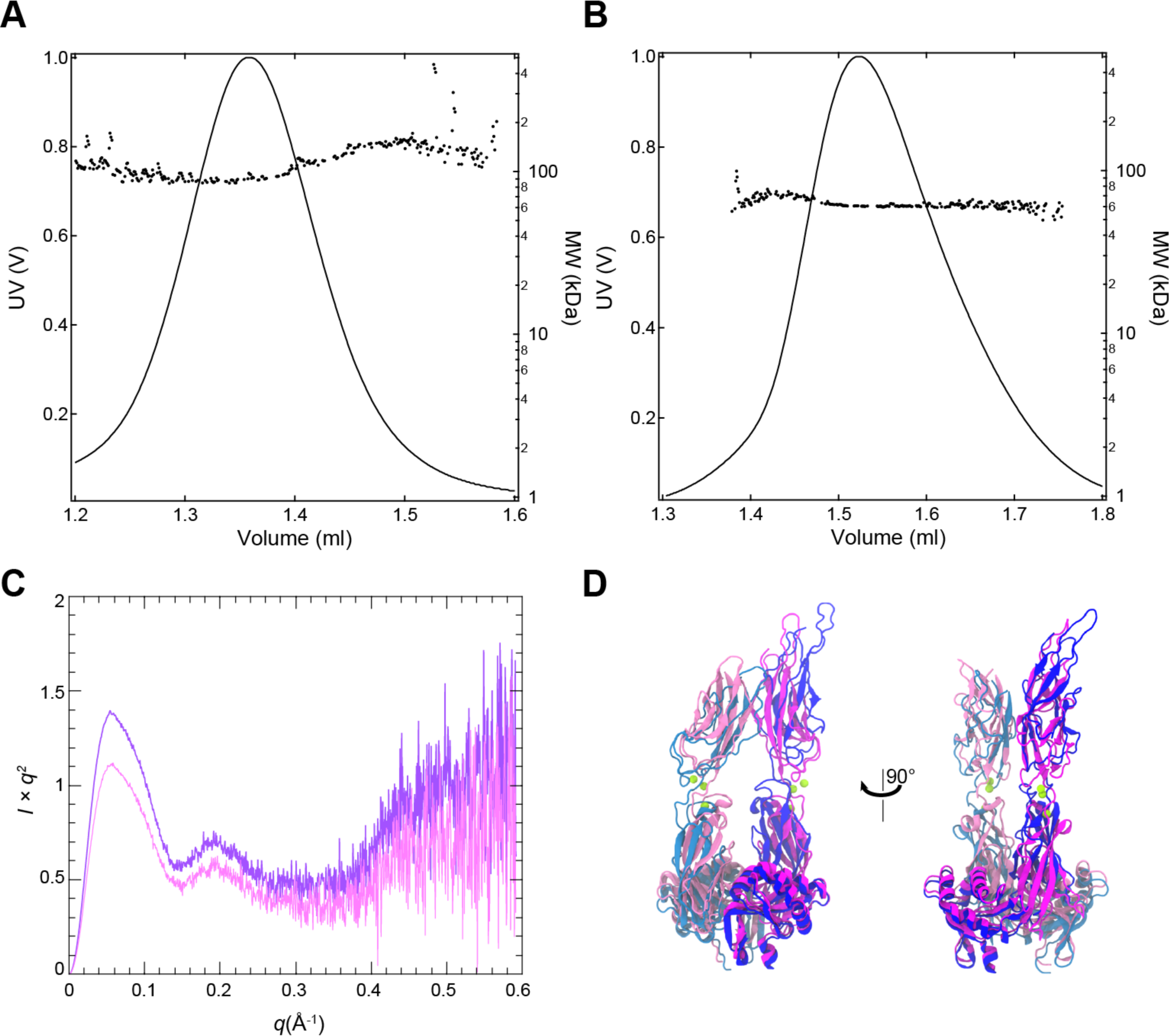
Dimerization of PCDH15 EC10-12 in solution. SEC-MALLS results for (***A***) *mm\** PCDH15 EC10-MAD12 and (***B***) *mm* PCDH15 EC10-MAD12. Solid line represents the UV signal measured at 280 nm, and dots represent the molecular weights calculated from the light scattering signal. (***C***) Kratky plot for *ss* and *mm* PCDH15 EC10-MAD12 (magenta and purple, respectively). The shape of the plot indicates the presence of folded protein with certain degree of flexibility for both samples. (***D***) Representative structure obtained for *ss* PCDH15 EC10-MAD12 from flexible refinement with SREFLEX (dark and light blue) superimposed to crystallographic model *ss* PCDH15 EC10-MAD12 (6BXZ, magenta and mauve).

**FIGURE S6.**
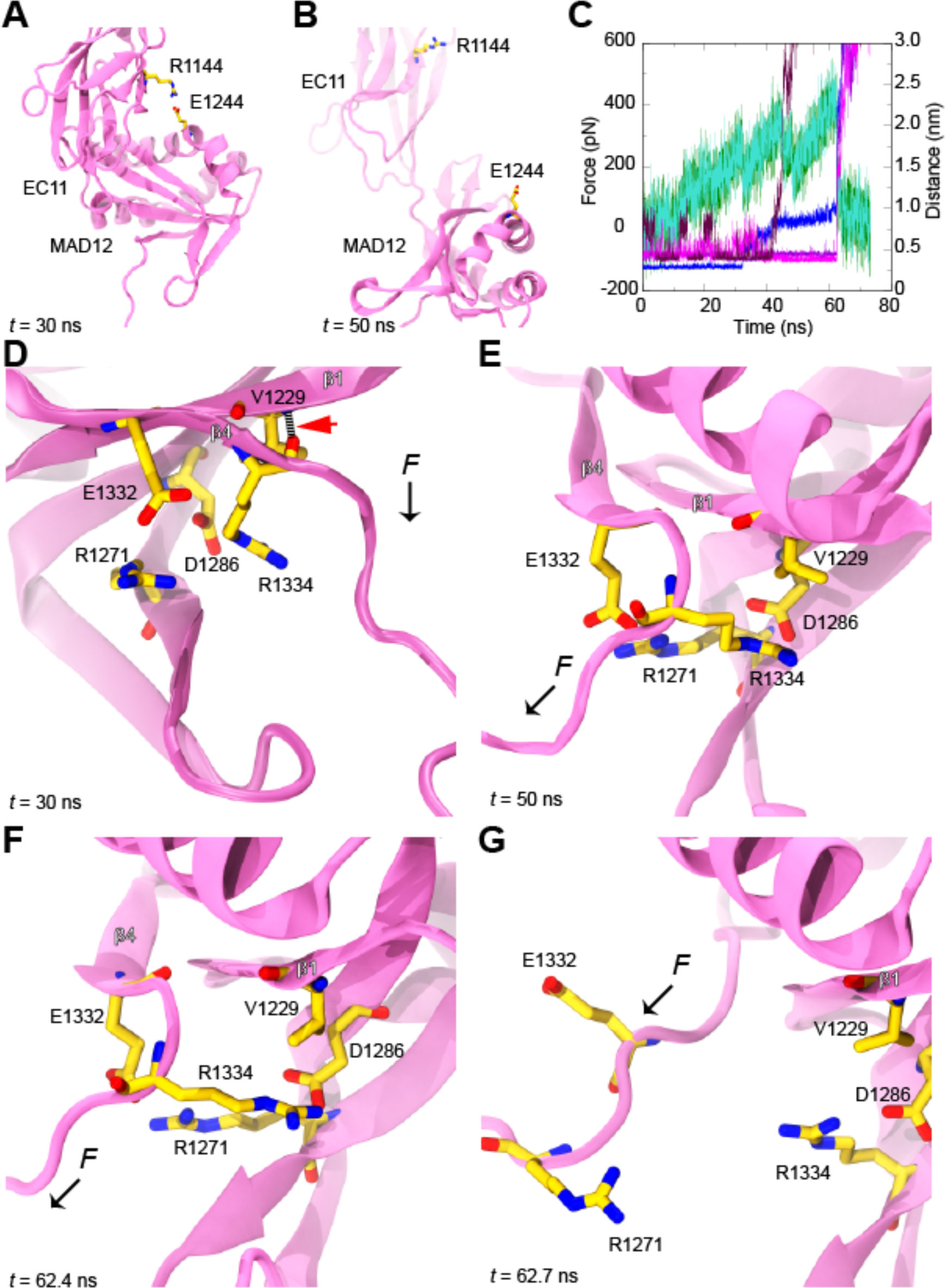
MAD12 unrolling and unfolding during stretching simulations of monomeric *ss* PCDH15 EC10-MAD12. (***A*-*B***) Snapshots of MAD12 before (*A*) and after (*B*) unrolling in simulation S1d. Unrolling induces MAD12’s hook swinging. (***C***) Force applied to *ss* PCDH15 EC9-MAD12 N-terminus (dark green) and C-terminus (turquoise) along with distance distances between atoms p.R1334-O and p.V1229-N (blue), p.R1334-Cζ and p.D1286-Cγ (violet), p.E1332-Cδ and p.R1271-Cζ (magenta), and p.E1244-Cδ and p.R1144-Cζ (maroon) versus time during simulation S1d. (***D***-***G***) Snapshots show unzipping of β4 and formation of electrostatic lock before unfolding at indicated time-points during simulation S1d. Red arrowhead indicates first backbone hydrogen bond ruptured during l4 unzipping. Black arrows indicate direction of applied force.

**FIGURE S7.**
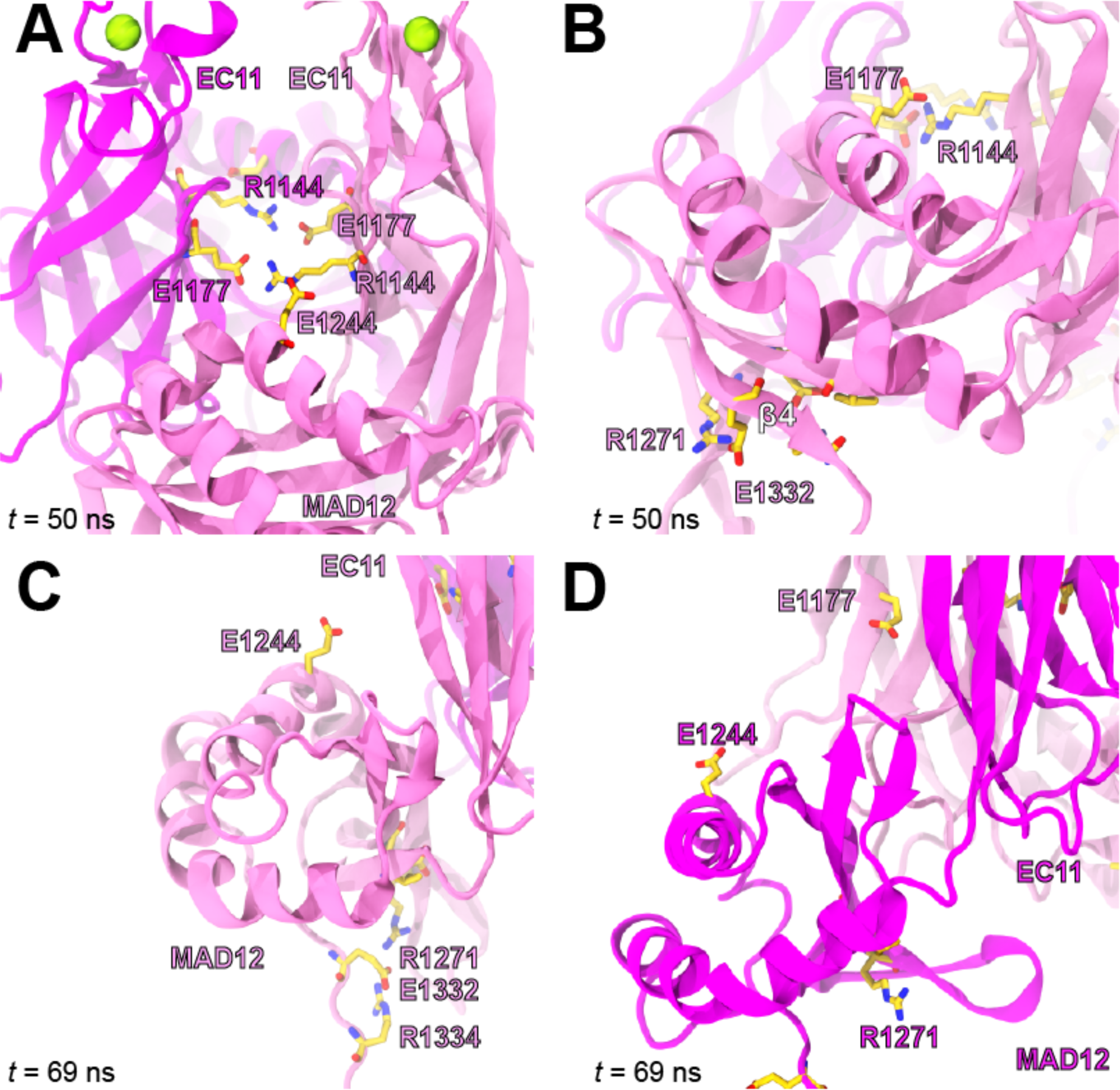
MAD12 unrolling and unfolding during stretching simulations of dimeric *ss* PCDH15 EC10-MAD12. (***A***-***D***) Snapshots of intermolecular interactions (*A*), key force-bearing residues during β4 unzipping (*B*), partial unrolling of MAD12 in one subunit (*C*), and complete unrolling of MAD12 in the other subunit (*D*) at indicated time points.

**FIGURE S8.**
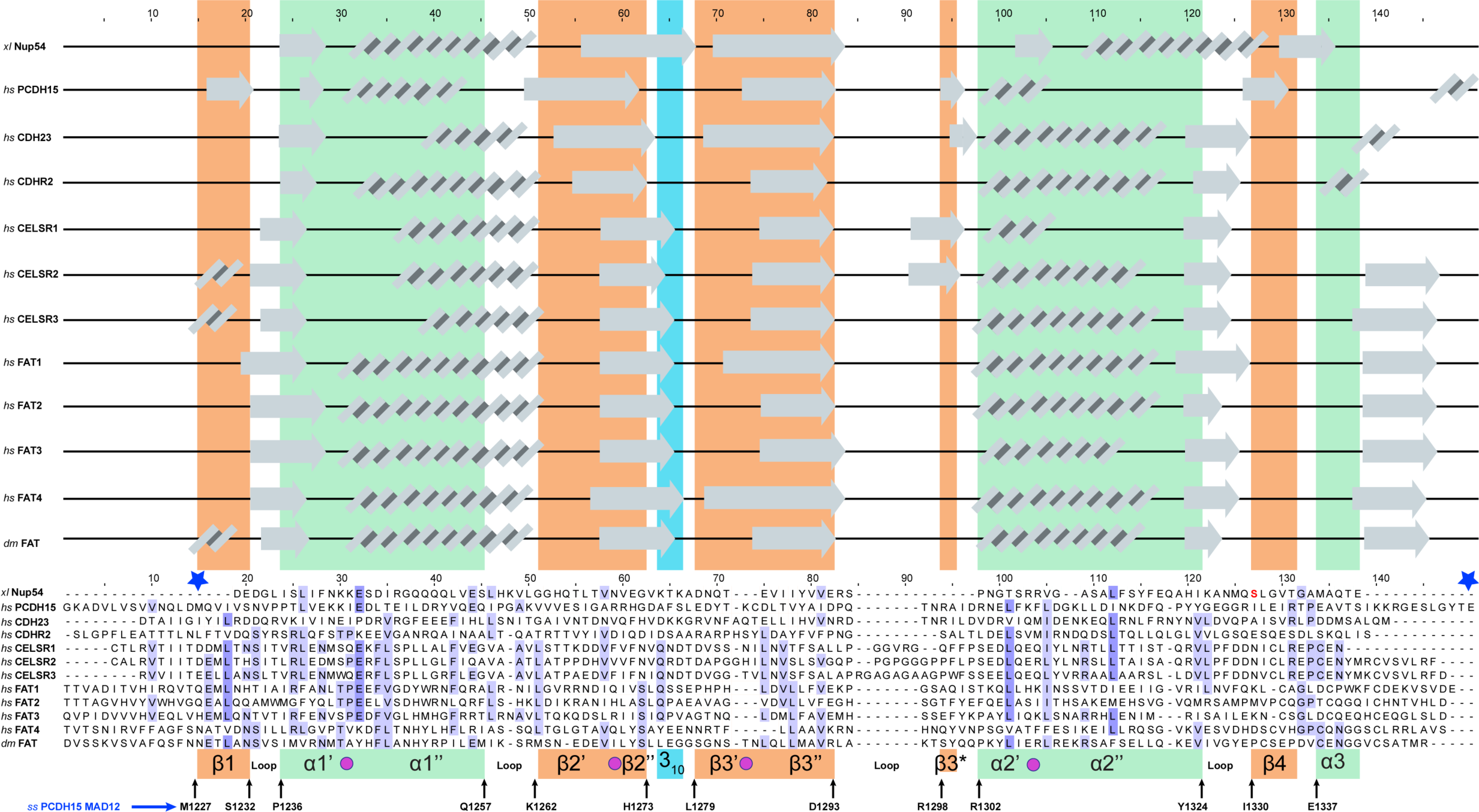
Multiple sequence alignments comparing MAD sequences from *xl* Nup54, *hs* PCDH15 MAD12, *hs* CDH23 MAD28, *hs* PCDH24 (CDHR2) MAD10, *hs* CELSR1-3 MAD10, *hs* FAT1-4 MADs, and *dm* FAT MAD35 (colored by % of sequence similarity using 1% threshold). White represents the lowest similarity, while blue represents the highest. Secondary structure from *ss* PCDH15 EC10-MAD12 is represented as colored columns in the background (β strands in green, α helices in orange, and 3_10_ in dark cyan). Predicted secondary structure (RaptorX) for each MAD are displayed on top of the sequence alignment. Secondary structure features from *ss* PCDH15 EC10-MAD12 and their starting and ending residues are labeled at the bottom of each colored column (residue numbering as in PDB ID 6BXZ). Blue stars represent the starting and ending residues of MAD12. Mutation p.127 (residue numbering from the alignment) in the *xl* Nup54 sequence is highlighted in red.

**FIGURE S9.**
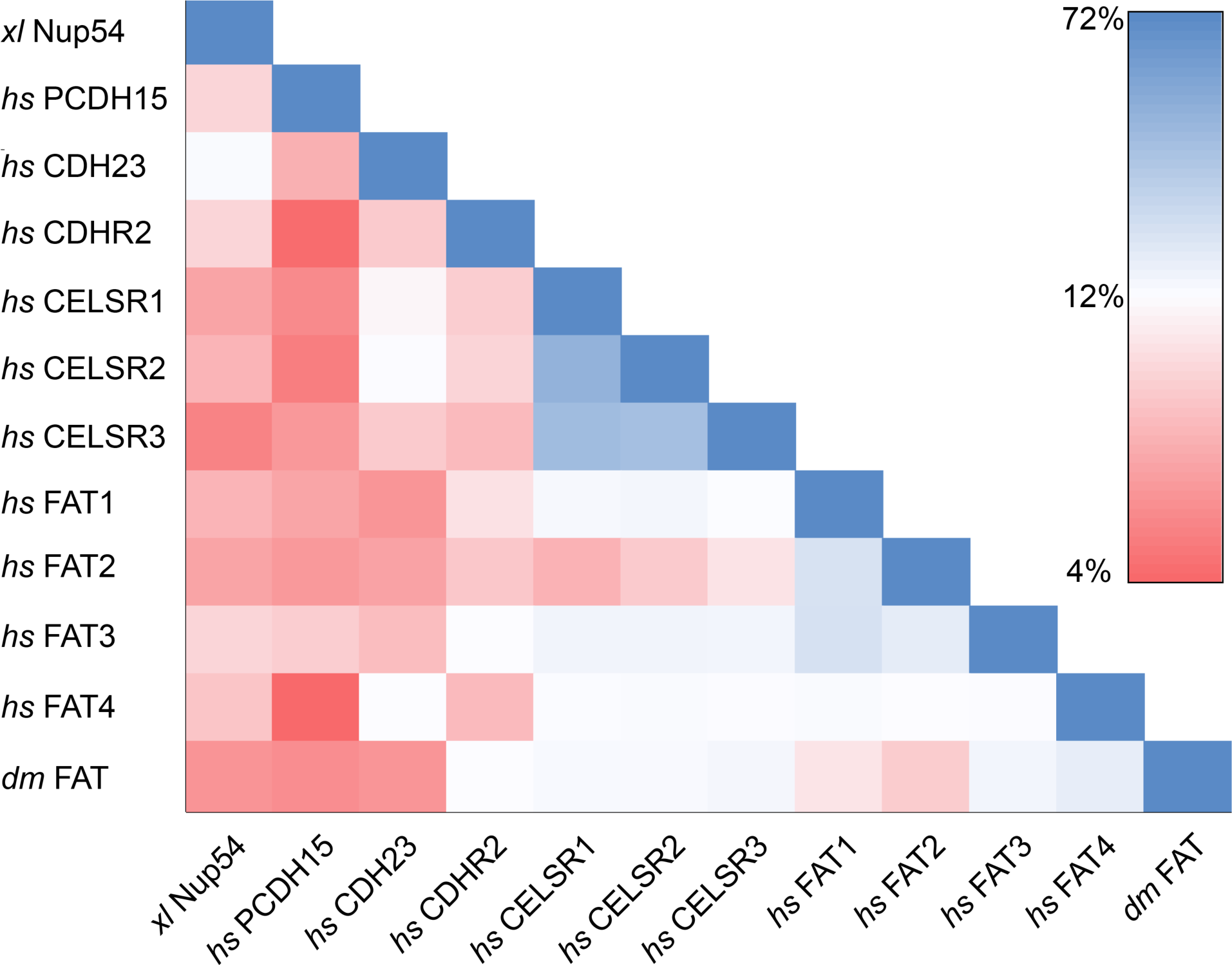
Heatmap of a sequence identity matrix among MADs. Sequence alignment from Figure S8 was 2 used as an input into the SIAS server to generate the matrix (89). Blue indicates the highest similarity, while red 3 indicates the lowest. Protein species are labeled as in Table S2. The average sequence identity among all MADs 4 was 15.64%.

**FIGURE S10.**
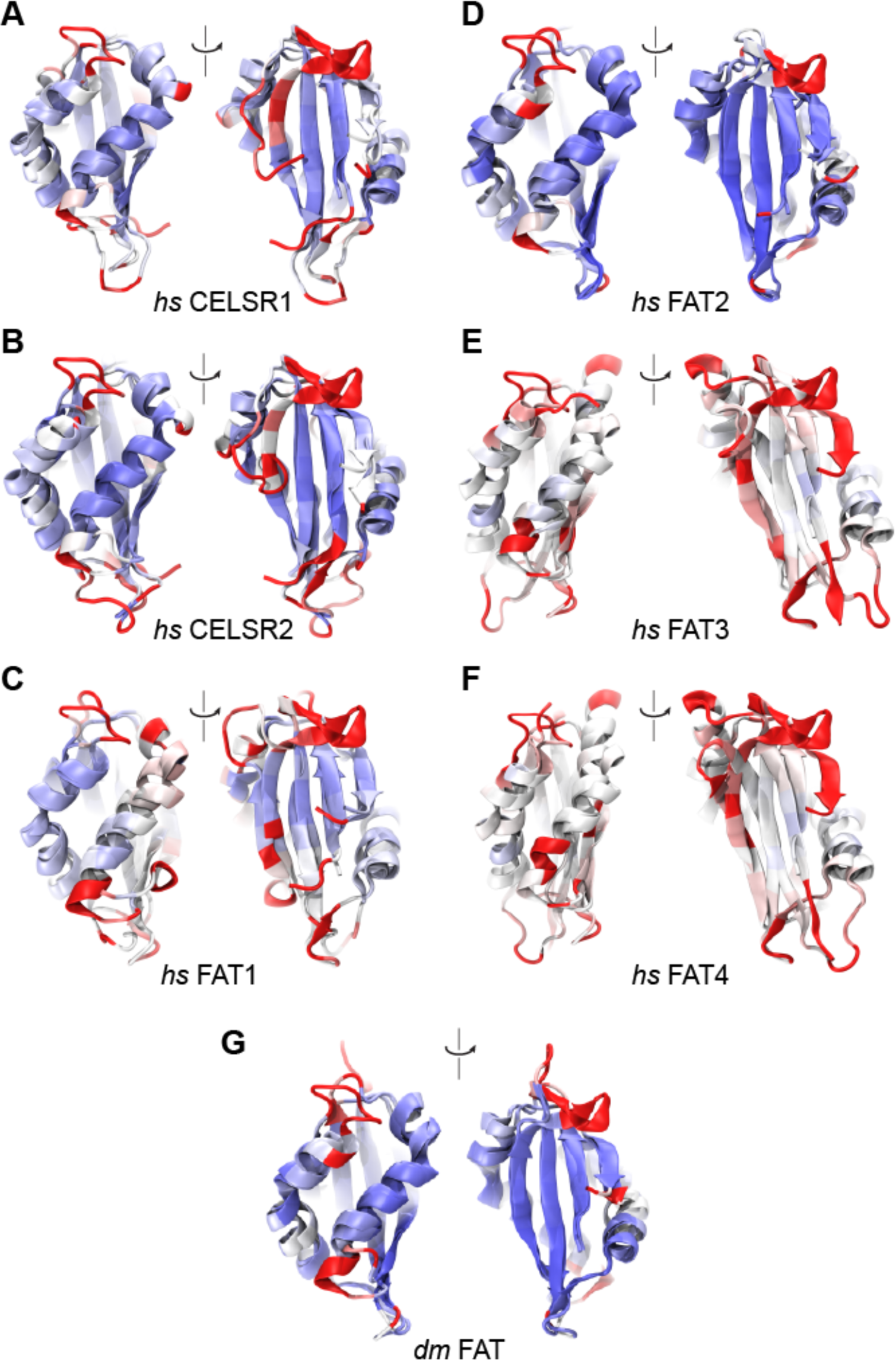
Structural models of MADs from atypical cadherins. (***A***-***G***) Front and back views of models superposed on the crystal structure of *ss* PCDH15 MAD12 (6BXZ) and shown as in Fig. 6 *B*.

**FIGURE S11.**
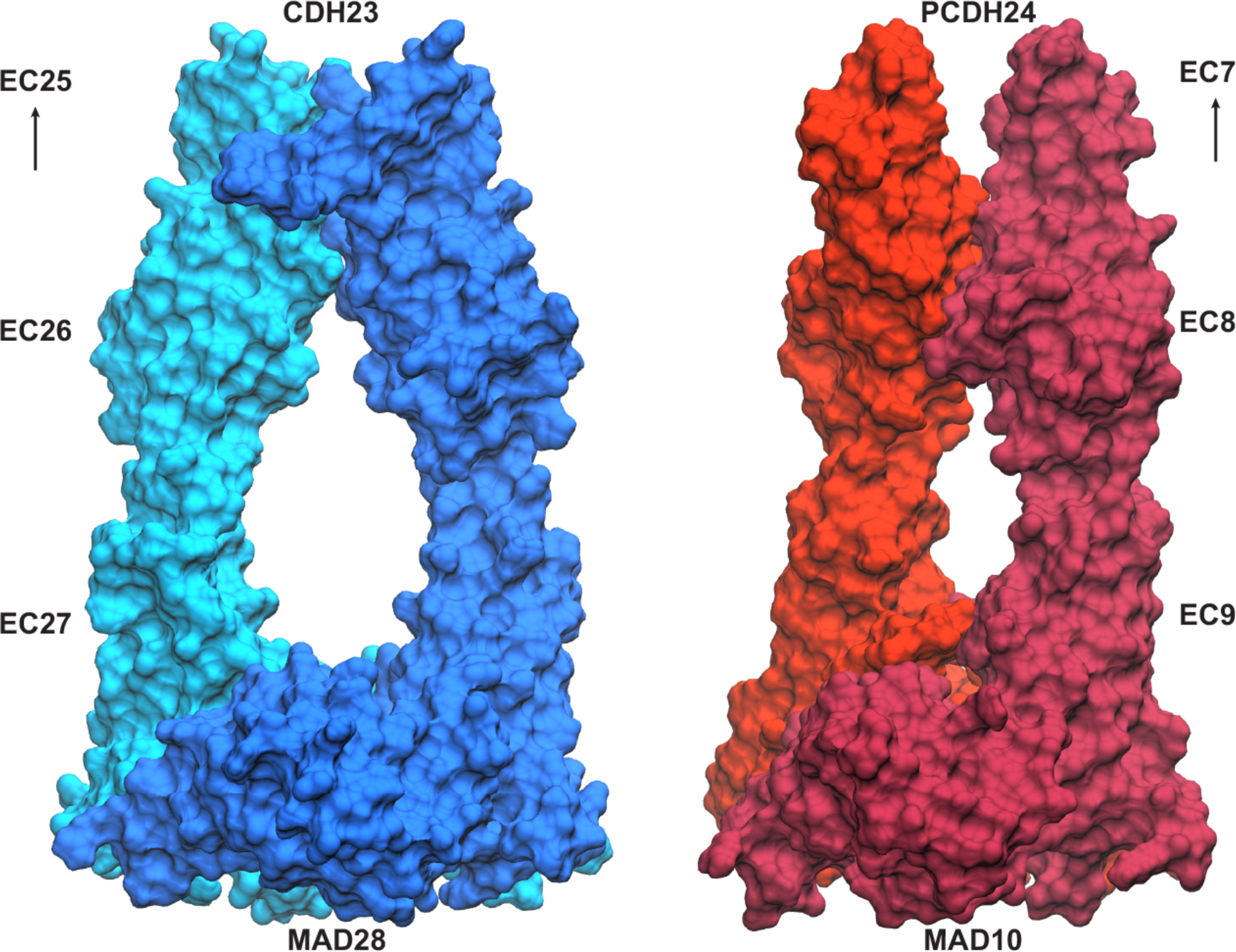
Models of *hs* CDH23 EC26-MAD28 (left) and *hs* PCDH24 EC8-MAD10 (right) homo dimers.

**TABLE S1.**
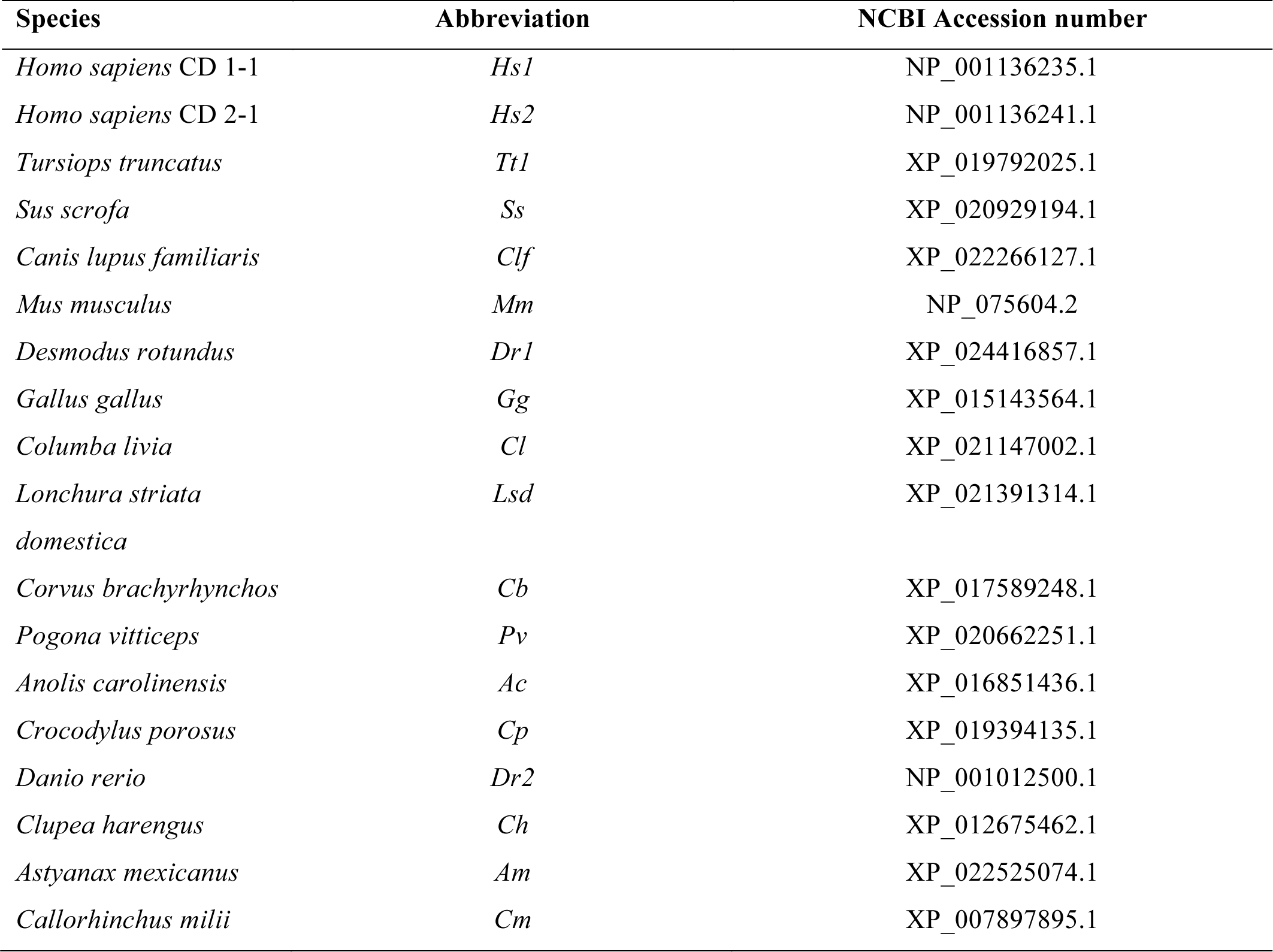
List of species and accession numbers used in multiple sequence alignment analysis of PCDH15 EC10-MAD12.

**TABLE S2.** List of proteins and accession numbers used in multiple sequence alignment of putative MADs.

| **Protein** | **Abbreviation** | **NCBI Accession number** | **Residue range** |
| --- | --- | --- | --- |
| *Xenopus laevis* Nup54 | *Xl* | XP_018090059.1 | 214 to 315 |
| *Homo sapiens* PCDH15 | *Hs* | NP_001136235.1 | 1239 to 1379 |
| *Homo sapiens* CDH23 | *Hs* | NP_071407.4 | 2942 to 3067 |
| *Homo sapiens* PCDH24 | *Hs* | NP_001165447.1 | 1027 to 1154 |
| *Homo sapiens* CELSR1 | *Hs* | NP_055061.1 | 1191 to 1314 |
| *Homo sapiens* CELSR2 | *Hs* | NP_001399.1 | 1113 to 1249 |
| *Homo sapiens* CELSR3 | *Hs* | NP_001398.2 | 1262 to 1397 |
| *Homo sapiens* FAT1 | *Hs* | XP_005262891.1 | 3622 to 3756 |
| *Homo sapiens* FAT2 | *Hs* | NP_001438.1 | 3610 to 3744 |
| *Homo sapiens* FAT3 | *Hs* | XP_016872668.1 | 3651 to 3783 |
| *Homo sapiens* FAT4 | *Hs* | NP_001278232.1 | 3694 to 3826 |
| *Drosophila melanogaster* FAT | *Dm* | NP_477497.1 | 3838 to 3971 |

**TABLE S3.**
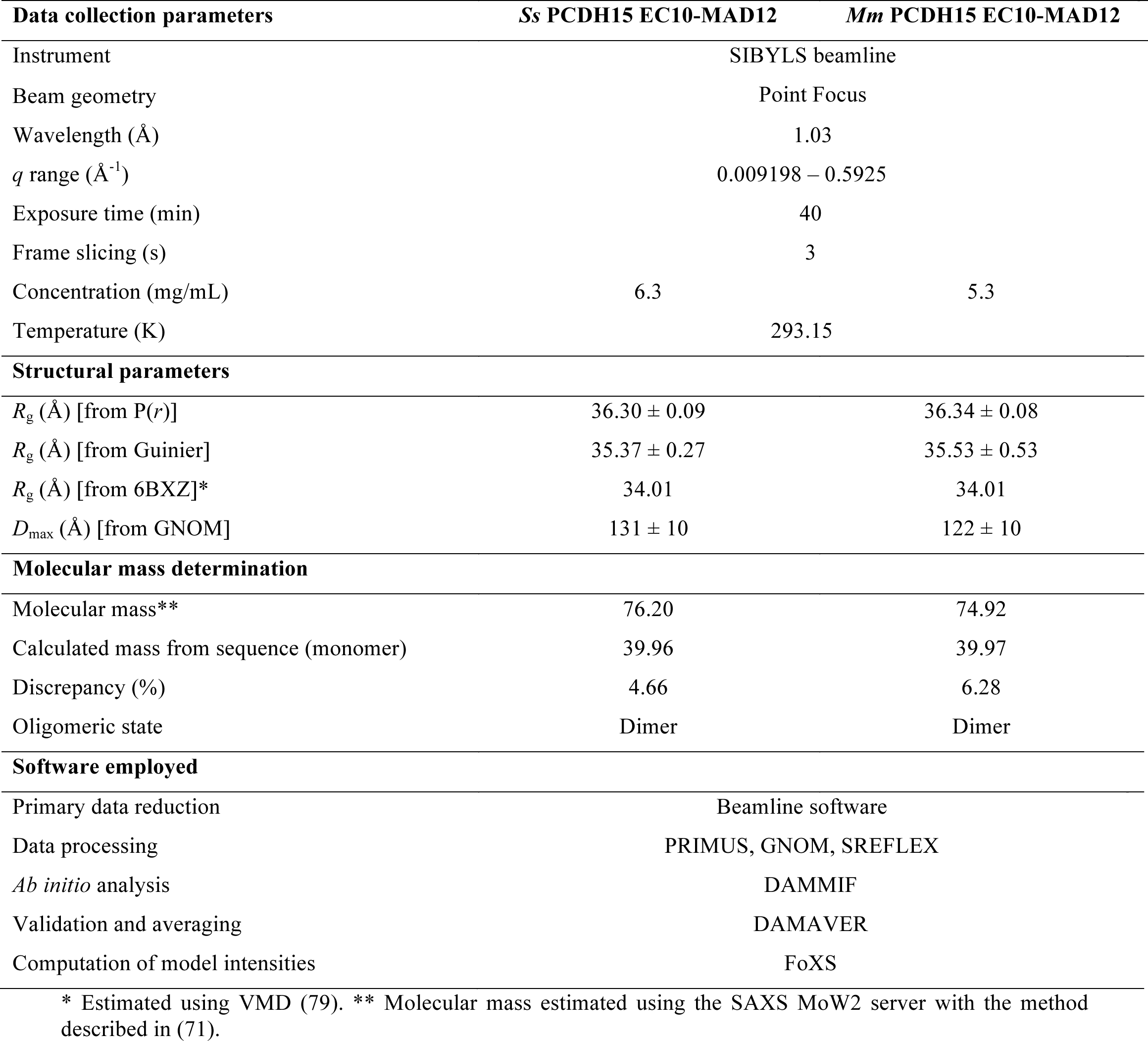
SAXS data collection and scattering-derived parameters

**TABLE S4.** Summary of SMD simulations.

| Label | System | $t_{\text{sim}}$ (ns) | Type | Start | Speed (nm/ns) | Average Peak Force (pN)<sup>b</sup> | Size (#atoms) | Initial Size (nm<sup>3</sup>) |
| --- | --- | --- | --- | --- | --- | --- | --- | --- |
| S1a | Monomer | 11.1 | EQ<sup>a</sup> | — | — | — | 211,618 | 27.3 × 9.7 × 8.4 |
| S1b |  | 3.0 | SMD<sup>c</sup> | S1a | 10 | 659.4 |  |  |
| S1c |  | 17.1 | SMD<sup>c</sup> | S1a | 1 | 529.3 |  |  |
| S1d |  | 73.2 | SMD<sup>c</sup> | S1a | 0.1 | 477.9 |  |  |
| S1e |  | 259.5 | SMD<sup>c</sup> | S1a | 0.02 | 313.7 |  |  |
| S2a | Dimer | 11.1 | EQ<sup>a</sup> | — | — | — | 279,489 | 27.1 × 10.8 × 10.1 |
| S2b |  | 3.0 | SMD<sup>c</sup> | S2a | 10 | 736.1 |  |  |
| S2c |  | 17.3 | SMD<sup>c</sup> | S2a | 1 | 571.4 |  |  |
| S2d |  | 68.9 | SMD<sup>c</sup> | S2a | 0.1 | 408.0 |  |  |
| S2e |  | 3.0 | SMD<sup>d</sup> | S2a | 10 | 1414.9 |  |  |
| S2f |  | 13.4 | SMD<sup>d</sup> | S2a | 1 | 1002.5 |  |  |
| S2g |  | 78.5 | SMD<sup>d</sup> | S2a | 0.1 | 721.0 |  |  |
| Total: |  | 559.1 |  |  |  |  |  |  |

<sup>a</sup> EQ indicates simulations that consisted of 1,000 steps of minimization, 100 ps of dynamics with the protein backbone constrained ( $k = 1$ kcal/mol/Å<sup>2</sup>), 1 ns of free dynamics in the $NpT$ ensemble ( $\gamma = 1$ ps<sup>-1</sup>), and 10 ns of free dynamics in the $NpT$ ensemble ( $\gamma = 0.1$ ps<sup>-1</sup>).
<sup>b</sup> Average peak force is calculated from the peak force measured on stretched Cα atoms or slabs (using 50-ps running averages).
<sup>c</sup> SMD simulation in which Cα atoms of residues p.G1009 and p.I1342 were attached to independent stretching springs.
<sup>d</sup> SMD simulation in which force was applied through slabs to protein ends (see Methods).

**TABLE S5.**
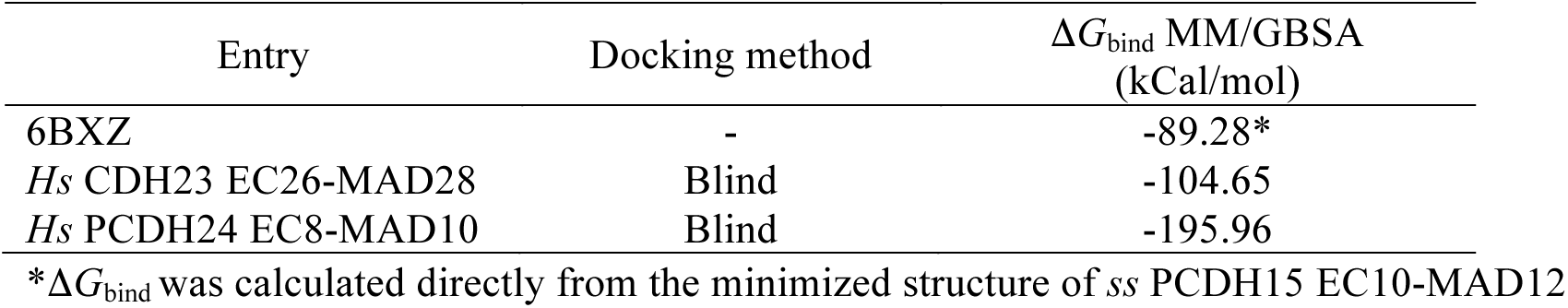
Predicted Δ*G*_bind_ values using Prime MM/GBSA

| Entry | Docking method | $\Delta G_{\text{bind}}$ MM/GBSA<br>(kCal/mol) |
| --- | --- | --- |
| 6BXZ | - | -89.28* |
| *Hs* CDH23 EC26-MAD28 | Blind | -104.65 |
| *Hs* PCDH24 EC8-MAD10 | Blind | -195.96 |

\* $\Delta G_{\text{bind}}$ was calculated directly from the minimized structure of *ss* PCDH15 EC10-MAD12

